# Kinetics of the immune response to *Eimeria maxima* in relatively resistant and susceptible chicken lines

**DOI:** 10.1101/757567

**Authors:** Abi Bremner, Sungwon Kim, Katrina Morris, Matthew J. Nolan, Dominika Borowska, Zhiguang Wu, Fiona Tomley, Damer P. Blake, Rachel Hawken, Pete Kaiser, Lonneke Vervelde

## Abstract

*Eimeria maxima* is a common cause of coccidiosis in chickens, a disease which has a huge economic impact on poultry production. Knowledge of immunity to *E. maxima* and the specific mechanisms that contribute to differing levels of resistance observed between chicken breeds and between congenic lines derived from a single breed of chickens is required. This study aimed to define differences in the kinetics of the immune response of two inbred lines of White Leghorn chickens that exhibit differential resistance (line C.B12) or susceptibility (line 15I) to infection by *E. maxima*. Line C.B12 and 15I chickens were infected with *E. maxima* and transcriptome analysis of infected jejunal tissue was carried out at 2, 4, 6 and 8 days post-infection (dpi). RNA-Seq analysis revealed differences in the rapidity and magnitude of cytokine transcription responses post-infection between the two lines. In particular, IFN-γ and IL-10 transcripts in the jejunum accumulated earlier in line C.B12 (at 4 dpi) compared to line 15I (at 6 dpi). Line C.B12 chickens exhibited increases of *IFNG* and *IL10* mRNA in the jejunum at 4 dpi, whereas in line 15I transcription was delayed but increased to a greater extent. RT-qPCR and ELISAs confirmed the results of the transcriptomic study. Higher serum IL-10 correlated strongly with higher *E. maxima* replication in line 15I compared to line C.B12 chickens. Overall, the findings suggest early induction of the IFN-γ and IL-10 responses, as well as immune-related genes at 4 dpi identified by RNA-Seq, may be key to resistance to *E. maxima*.

## Introduction

Coccidiosis, which in poultry is caused by apicomplexan parasites of the genus *Eimeria*, causes huge economic losses to the global poultry industry through decreased feed efficiency, reduced weight gain, increased mortality, and the cost of prophylaxis and therapy. It is the most economically important parasitic condition of poultry (1, 2). One of seven *Eimeria* species that can infect chickens, *Eimeria maxima* is commonly diagnosed in commercial chicken flocks (3, 4) and specifically invades and parasitizes enterocytes of the jejunum where it can cause pathological lesions, resulting in villus destruction and malabsorptive disease symptoms (5). Currently, control of *Eimeria* is primarily achieved through in-feed prophylaxis with anti-coccidial drugs or by vaccination with live, or live-attenuated parasites. However, resistance to anticoccidial drugs is common (6) and vaccination is complex, requiring the preparation and administration of admixtures of between three and eight different lines of parasite to confer adequate protection against filed challenge (7). A potential alternative method of control could be to selectively breed chickens that have enhanced resistance to *Eimeria*; however, this requires knowledge of the natural host immune response to *Eimeria* and the identification of biomarkers of resistance.

Understanding the immunological basis of resistance to *Eimeria* is an important step towards identifying biomarkers of resistance for the selection of relatively resistant individuals within commercial breeding stocks. Inbred lines 15I (MHC type B^15^) and C.B12 (MHC type B^12^) are White Leghorn chickens which display differential resistance and susceptibility to *E. maxima* based on oocyst output. Following primary infection line C.B12 chickens shed fewer oocysts compared to line 15I, but both lines display complete immune protection against homologous secondary infection after which no oocysts are produced (8, 9). Additionally, two-fold higher levels of *E. maxima* DNA have been detected in the intestinal tissue of line 15I compared to line C.B12 chickens at 5 days post-infection (dpi) (10). Another study reported that line FP (MHC type B^15^/B^21^) chickens produce more oocysts than line SC (MHC type B^2^) chickens after infection with *E. maxima* (11). Although these chicken lines were bred for specific MHC types, the immunological basis underlying resistance and susceptibility to *E. maxima* is not well characterized.

Following *E. maxima* infection, cell-mediated immunity and host genetic variation in T-cell responses appear to be central to the induction of protective immunity (8, 12). Although parasite-specific antibodies can protect against *E. maxima* infection, (13, 14, 15), bursectomised (B-cell deficient) chickens were no more susceptible to *E. maxima* challenge than non-bursectomised control birds (16), suggesting that antibodies are not necessary for elimination of the parasite. An array of cell-mediated responses are a prominent feature of coccidiosis and attempts have been made to correlate these responses with immunity. Primary *E. maxima* infection leads to an increased percentage of CD8 and γδ T cells in peripheral blood leukocytes (PBL) in relatively resistant (line C) chickens compared to relatively susceptible (line 15I and 6_1_) chickens, whereas there was no significant difference in CD4 and αβ2 T cells between these lines of chickens (8). On the other hand, increased numbers of CD4 lamina propria lymphocytes (LPL), but not intraepithelial lymphocytes (IEL), were observed in relatively susceptible Light Sussex chickens at 3 dpi (17), while CD8 LPL and IEL were increased at 4 dpi (18). Overall, there were more CD8 than CD4 cells within the gut during *E. maxima* infection (8, 18). During *E. maxima* infection, significantly increased γδ and αβ1 T cells were reported in the epithelium at later time points (11 dpi), while αβ2 T cells in the lamina propria increased at 4 and 11 dpi (18, 19), although there was induction time variation dependent on the genetic background of the chickens and the nature of the challenge dose.

Interferon (IFN)-γ, a key signature cytokine of Th1-controlled immune responses, is a major cytokine mediating a protective immune response against many intracellular pathogens including viruses (20, 21), *Salmonella* spp. (22) and *Eimeria* spp. (23, 24). Early studies showed that increased serum IFN-γ protein and gut *IFNG* mRNA levels are strongly associated with *E. acervulina* (24, 25), *E. maxima* (15) and *E. tenella* (26) infection. During *E. maxima* infection, significantly increased IFN-γ protein was observed in both the gut and serum of relatively susceptible (line SC) chickens, and serum IFN-γ levels are positively correlated with faecal oocyst shedding (15). Additionally, *E. maxima* infection leads to induction of *IFNG* mRNA levels in the IEL population of relatively susceptible (line SC) chickens during primary infection, but not secondary infection (19), implicating the importance of IFN-γ in the cellular immune response to primary *Eimeria* spp. infection.

Interleukin (IL)-10 is an anti-inflammatory and regulatory cytokine and is important in balancing inflammatory responses to pathogens. During the characterization of biological roles of chicken IL-10, its potential role as a biomarker for *Eimeria* spp. infection was suggested. Increased *IL10* mRNA levels were observed in the spleen and the small intestine of relatively susceptible (line 15I) chickens during *E. maxima* infection compared to non-infected chickens, but not in relatively resistant (line C.B12) chickens (27). Moreover, uninfected relatively susceptible chickens had significantly higher *IL10* mRNA levels in the spleen compared to relatively resistant chickens (27), suggesting that levels of constitutive IL-10 expression may be dependent on host genetics. Further studies showed increased *IL10* mRNA levels in the liver and caecum (28) and IL-10 protein in the serum (29) during *E. tenella* infection. Furthermore, antibody-mediated depletion of luminal IL-10 reduced oocyst shedding in broilers given an attenuated *Eimeria* spp. vaccine (30).

The present study aimed to characterise in detail the kinetics of the immune responses of relatively resistant (line C.B12) and susceptible (line 15I) inbred chickens to *E. maxima* infection. To identify phenotypes that associate with resistance to *E. maxima*, we investigated differences in gene expression and the systemic and local kinetics of the IFN-γ, and IL-10 response between the two lines. Transcriptomic analysis revealed that interferon-mediated immune responses were induced in line C.B12 chickens at 4 dpi compared to the relatively susceptible line 15I chickens at 6 dpi. Both *IFNG* and *IL10* were expressed in similar patterns during the course of infection in each line. Line C.B12 chickens produced higher levels of IFN-γ and IL-10 proteins in the jejunum and serum until 5 dpi compared to line 15I chickens, whereas by 6-8 dpi line 15I chickens produced higher levels of both.

We also found that IFN-γ and IL-10 protein expression and mRNA transcription was highly correlated with parasite burden, with the strongest correlation between parasitaemia and serum IL-10 in line 15I chickens.

## Results

### Comparison of body weight gain and E. maxima load between relatively resistant and susceptible chickens

To examine the impact of *E. maxima* infection on the growth of line C.B12 (relatively resistant) and line 15I (relatively susceptible) chickens, the percentage weight gains were calculated for individual birds from 2 days prior to *E. maxima* infection to the time of culling (Figure 1A). *E. maxima* infection did not affect body weight gain (BWG) compared to control birds and there was no difference between the two lines during the course of the experiment. The low challenge dose did not result in lesions in the gut of either line.

**Figure 1.**
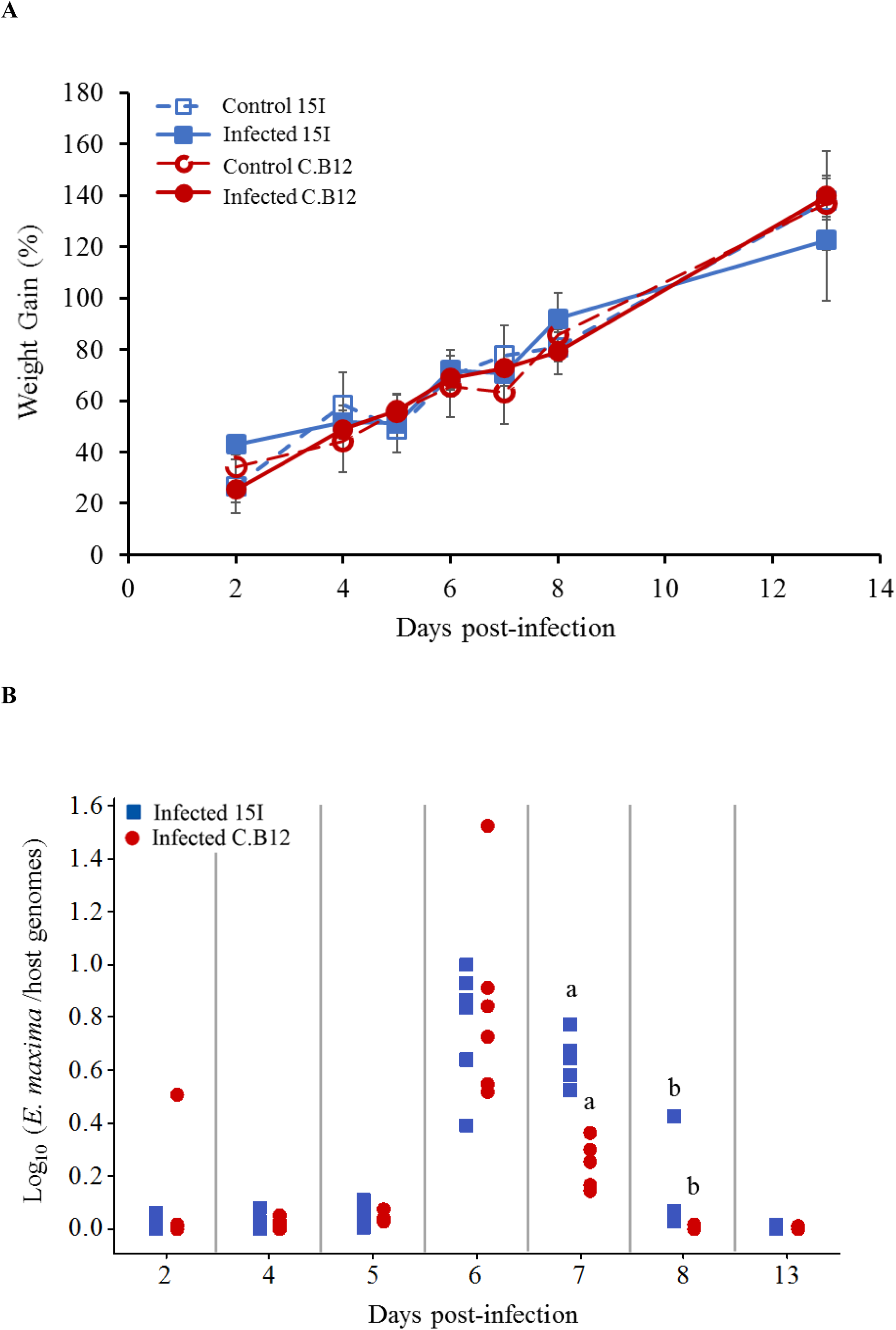
Body weight gains and parasite replication in line C.B12 and 15I chickens following *E. maxima* infection. Three-week-old birds were orally infected with 100 sporulated *E. maxima* oocysts (*n* = 5 per line) or sterile water (*n* = 3 per line). (A) Percentage of body weight gains were calculated for individual birds from 2 days prior to inoculation to time of culling at time points as indicated. The results were presented as the mean percentage of body weight gain and error bars represent standard deviation. (B) *Eimeria maxima* replication was quantified by qPCR targeting the MIC1 gene. The results were presented as the ratios of parasite genome vs host genome copy numbers for individual birds. Matching letters indicate significant differences between the two lines at *p* < 0.05 on the same day (*n* = 5 per time point).

*E. maxima* genome copy numbers sharply increased at 6 dpi to similar levels in both lines of birds (Figure 1B). Thereafter the genome copy numbers decreased in both lines but was significantly higher in the jejunum of line 15I compared with line C.B12 chickens at 7 and 8 dpi. By 13 dpi, no difference in *E. maxima* genome copy number was apparent between the two lines. *Eimeria* genomes remained detectable one day later in relatively susceptible line 15I chickens.

### Comparison of global kinetic gene expression profiles between relatively resistant and susceptible chickens during E. maxima infection

To explore host responses to *E. maxima* infection and the genetics underlying the relative differences in resistance and susceptibility between line C.B12 and 15I chickens, transcriptome analysis was performed. Differentially expressed genes (DEGs) were identified within the jejunum anterior to Meckel’s diverticulum, site of peak *E. maxima* replication, between control and infected chickens of each line at 2, 4, 6 and 8 dpi under the following conditions: False Discovery Ratio (FDR) < 0.05 and log(Fold Change (FC)) > 1.6 (Table 1: Table S1 and S2). Line 15I chickens showed very little response at 2 dpi (5 DEGs) and 4 dpi (3 DEGs), but had a large number of DEGs at 6 dpi (1124 DEGs). In contrast, line C.B12 had already established a substantial response by 4 dpi (177 DEGs), but also demonstrated a peak response at 6 dpi (666 DEGs). In line C.B12, 42.2% and 26.8% of the DEGs were immune-related in function at 4 and 6 dpi, respectively. In line 15I, there was no differential expression in immune-related genes at 4 dpi, while 29.2% of DEGs at 6 dpi were immune-related. Immune genes upregulated strongly in both lines at day 6 included *IFNG*, chemokines and complement components. Analysis using the Markov clustering algorithm indicated that samples from line C.B12 at 6 dpi and line 15I at 8 dpi were the furthest distance from controls, indicating that globally the peak responses may occur at these times (Figure 2A). A network graph of unbiased gene-to-gene clustering was constructed (Figure 2B and Table S3). Out of 12 clusters, cluster 5 revealed a set of 163 genes (Figure 2C), which included *IFNG* and *IL10*, that were strongly elevated at 6 dpi in both lines of chickens, but also earlier at 4 dpi in line C.B12 chickens. Further functional analysis revealed that genes of this cluster are mainly involved in interferon signalling, the Th1 pathway, and the Th1 and Th2 activation pathways. Genes in Cluster 5 included the IFN-α/β receptor (*IFNAR*), *IFNG*, interferon regulatory factor (*IRF*), protein tyrosine phosphatase (*PTPN2*), suppressor of cytokine signalling 1 and 2 (*STAT1* and *STAT2*), transporter 1 ATP binding cassette (*TAP1*; participates in the interferon signalling pathway), *CD80*, *CD274*, delta like canonical Notch ligand 4 (*DLL4*), *IL10*, *IL12A* (participates in the Th1 pathway and Th1 and Th2 activation pathways), C-C motif chemokine ligand 1 (*CCL1*), *CCL4*, complement components 1s (*C1S*), *C1R* and genes involved in the JAK-STAT cascade.

**Figure 2.**
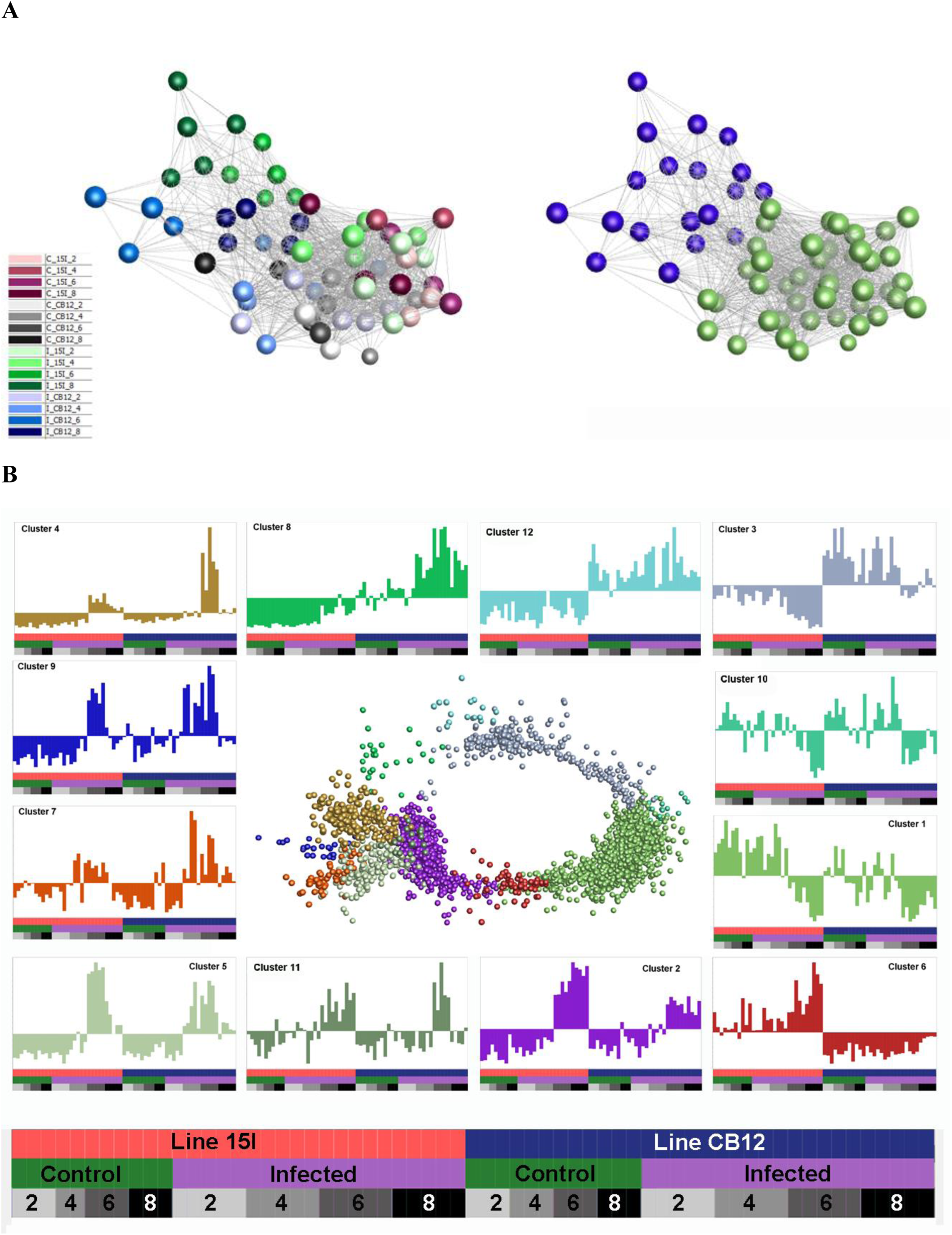
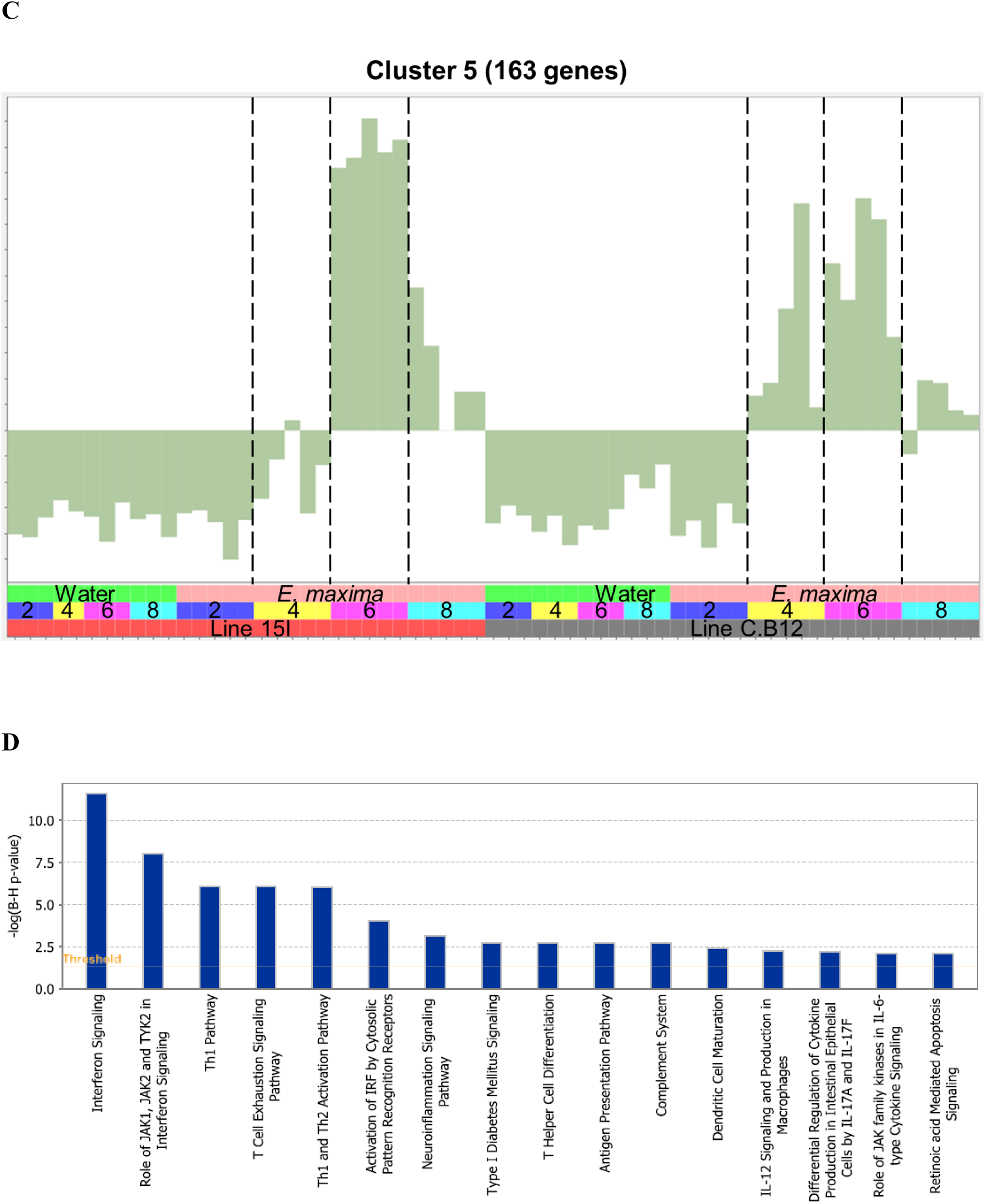
A network graph of unbiased sample-to-sample and gene-to-gene clustering. The sample-sample network shown on the left-hand side is coloured based on treatment group, while the right-hand network is coloured by Markov clustering of samples (A). Gene-gene network graph of Markov clustered genes (B), includes normalised expression across samples (mean-centered scaling) of each cluster in the surrounding charts. Genes in cluster 5 (C) including IFN-γ and IL-10 have strongly elevated expression at 6 dpi in both lines of chickens, but also earlier at 4 dpi in line C.B12 chickens. Pathways enriched in cluster 5 are shown in (D).

**Table 1.** Number of differentially expressed genes (DEGs)

| Contrast | Total number of DEGs<sup>1</sup> |  | Immune-related genes (%)<sup>2</sup> |  |
| --- | --- | --- | --- | --- |
|  | Up | Down | Up | Down |
| Line 15I infected vs. control at 2 dpi | 3 | 2 | 0 | 0 |
| Line 15I infected vs. control at 4 dpi | 1 | 2 | 0 | 0 |
| Line 15I infected vs. control at 6 dpi | 845 | 279 | 24.7 | 4.5 |
| Line 15I infected vs. control at 8 dpi | 303 | 335 | 14.8 | 6.6 |
| Line C.B12 infected vs. control at 2 dpi | 12 | 1 | 0 | 100 |
| Line C.B12 infected vs. control at 4 dpi | 68 | 109 | 42.2 | 0 |
| Line C.B12 infected vs. control at 6 dpi | 314 | 352 | 17.2 | 9.6 |
| Line C.B12 infected vs. control at 8 dpi | 48 | 123 | 29.6 | 9.6 |

<sup>1</sup>The threshold: FDR < 0.05 and abs FC > 1.6.
<sup>2</sup>Immune-related genes are the percentage of genes with human orthologs in which the human ortholog is categorized as having immune function.

### Kinetics of differential gene expression in relatively resistant and susceptible chickens

At 2 and 4 dpi, *E. maxima*-infected line 15I chickens had only 5 and 3 significant DEGs respectively, compared to control birds, although none of these were immune-related (Table 1). At 6 dpi, the largest increase in the expression of immune-related genes in line 15I was observed with 25% of upregulated genes with known functions having immune roles (Table 1). The pathways associated with the response of line 15I chickens at 6 dpi were primarily involved in T cell differentiation including differentiation into Th1 and Th2 subsets (Figure 3A). Gene ontology (GO) term enrichment analysis also highlighted the IL-21, IL-2 and IFN-γ pathways (Table S4). The highest upregulated protein coding genes were a complement receptor (homolog of *CR1*), *IFNG* and a gene involved in lipid metabolism (*ELOVL3*). Significant upregulation of immune-related genes were still observed at 8 dpi in line 15I, with 14.8% of 638 DEGs being immune-related. Upregulated genes at 8 dpi are involved in the complement and cell replication pathways, while genes associated with coagulation were downregulated (Figure 3B, Table S4). *IFNG* and *IL10* continued to be significantly upregulated at 8 dpi, and chemokines *CCL26* and *chCCLi7* were highly upregulated.

**Figure 3:**
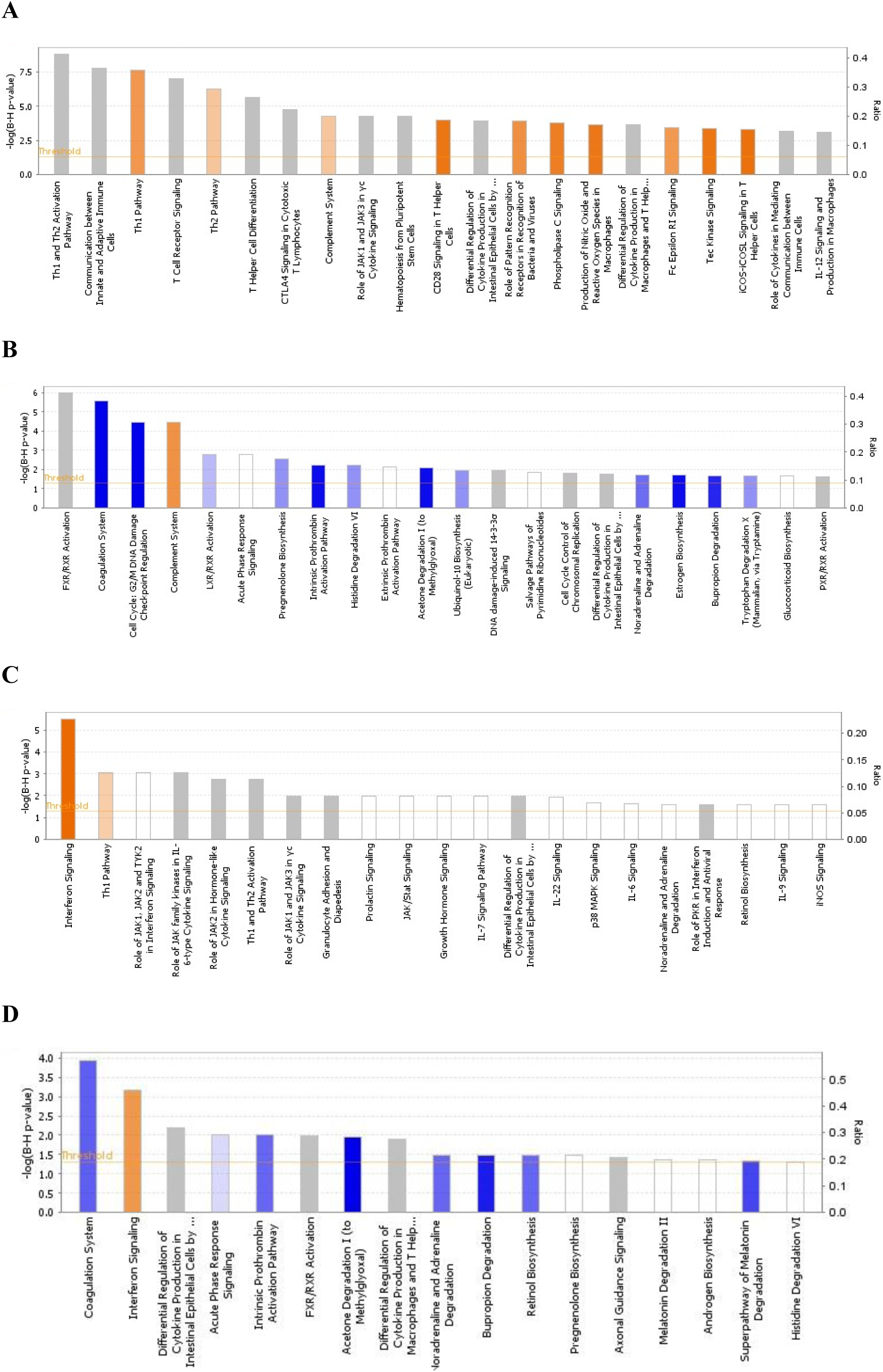
Ingenuity pathway analysis during *E. maxima* infection in line 15I at 6 (A) and 8 (B) dpi and line C.B12 chickens at 4 (C) and 6 (D) dpi. Colour based on Z-score with orange indicating activated pathways and blue indicating de-activated pathways.

In comparison with relatively susceptible (line 15I) chickens, relatively resistant (line C.B12) chickens developed immune responses to *E. maxima* infection as early as 2 and 4 dpi. A total of 13 DEGs were identified between *E. maxima*-infected line C.B12 compared to non-infected chickens at 2 dpi (Table 1). Of these genes, most of those which were upregulated were associated with erythrocytes. At 4 dpi in line C.B12, 42% of 177 DEGs were immune-related genes (Table 1).

Further functional analysis revealed that genes involved in the interferon signalling and Th1 pathways were strongly upregulated (Figure 3C) including: interferon-induced protein with tetratricopeptide repeats 1 (*IFIT1*), MX dynamin GTPase 1 (*MX1*, participates in the interferon signalling pathway), *CD274*, suppressor of cytokine signalling 3 (*SOCS3*, participates in Th1 pathway), *IFNG*, *SOCS1* and signal transducer and activator of transcription 1 (*STAT1*, participates in both pathways). GO term enrichment analysis indicated genes associated with T cell activity, the IFN-γ pathway, the JAK-STAT cascade and response to virus were strongly upregulated (Table S5). The highest upregulated protein coding genes were *IFNG*, a homolog of lyzosyme-G (ENSGALG00000044778), *CCL4* and GTPase, very large interferon inducible pseudogene 1 (*GVINP1*). Some of the upregulated interferon-stimulated genes such as radical S-adenosyl methionine domain containing 2 (*RSAD2*), *IFIT1*, *MX1* and 2’-5’-oligoadenylate synthetase like (*OASL*) were not significantly upregulated at any time point in line 15I chickens (Table S1). At 6 dpi, interferon and T-cell related genes continued to be upregulated in line C.B12, with the highest peak of *IFNG* and *IL10* expression observed (Figure 3D, Table S2). By 8 dpi, the response of line C.B12 chickens had subsided with only 172 DEGs (Table 1). These genes were varied and no significantly enriched GO terms were identified. Ingenuity pathway analysis revealed that only the coagulation pathway – regulated by fibrinogen gamma (*FGG*), kininogen 1 (*KNG1*), plasminogen (*PLG*) – was significantly downregulated in line C.B12 at 8 days post *E. maxima* infection.

### Comparison of the immune responses between line C.B12 and line 15I chickens

To directly compare the response to infection in the two chicken lines, the DEGs with the highest mean difference in logFC during *E. maxima* infection between the lines were examined, and the top 50 were plotted in a heatmap (Figure 4A). DEGs uniquely upregulated in line 15I included cytokines and genes associated with chemotaxis (TNF receptor superfamily member 13C (*TNFRSF13C*), C-X-C motif chemokine ligand 13 (*CXCL13*), chemokine ah221 (*CCL9*) and Pre-B lymphocyte protein 3 (*VPREB3*). A group of interferon-stimulated viral response genes (*IFIT5*, *RSAD2*, *MX1*, *OASL* and ubiquitin specific peptidase 18 (*USP18*) were upregulated at 4 and 6 dpi in line C.B12 but not line 15I chickens, further highlighting that this pathway is responding at a relatively higher level in line C.B12 compared to line 15I chickens. An additional group of genes was strongly upregulated in line C.B12 at 6 dpi only. Many of these genes are involved in epidermis development (keratin 75 (*KRT75*), *KRT15*, *KRT12*, ALX homeobox 4 (*ALX4*), homeobox B13 (*HOXB13*), suggesting that tissue repair is occurring at this time point in line C.B12, but may be delayed in line 15I.

**Figure 4.**
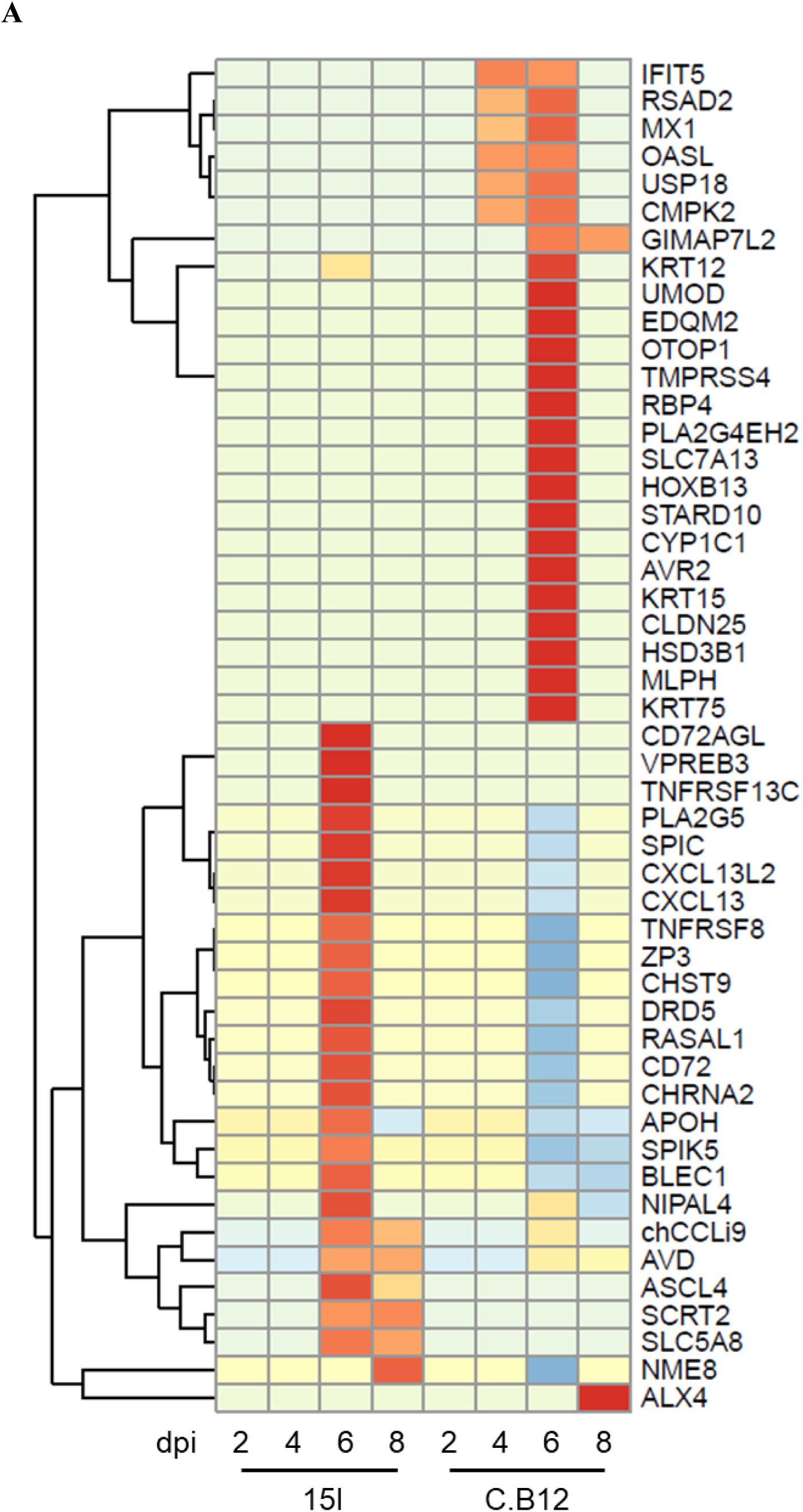
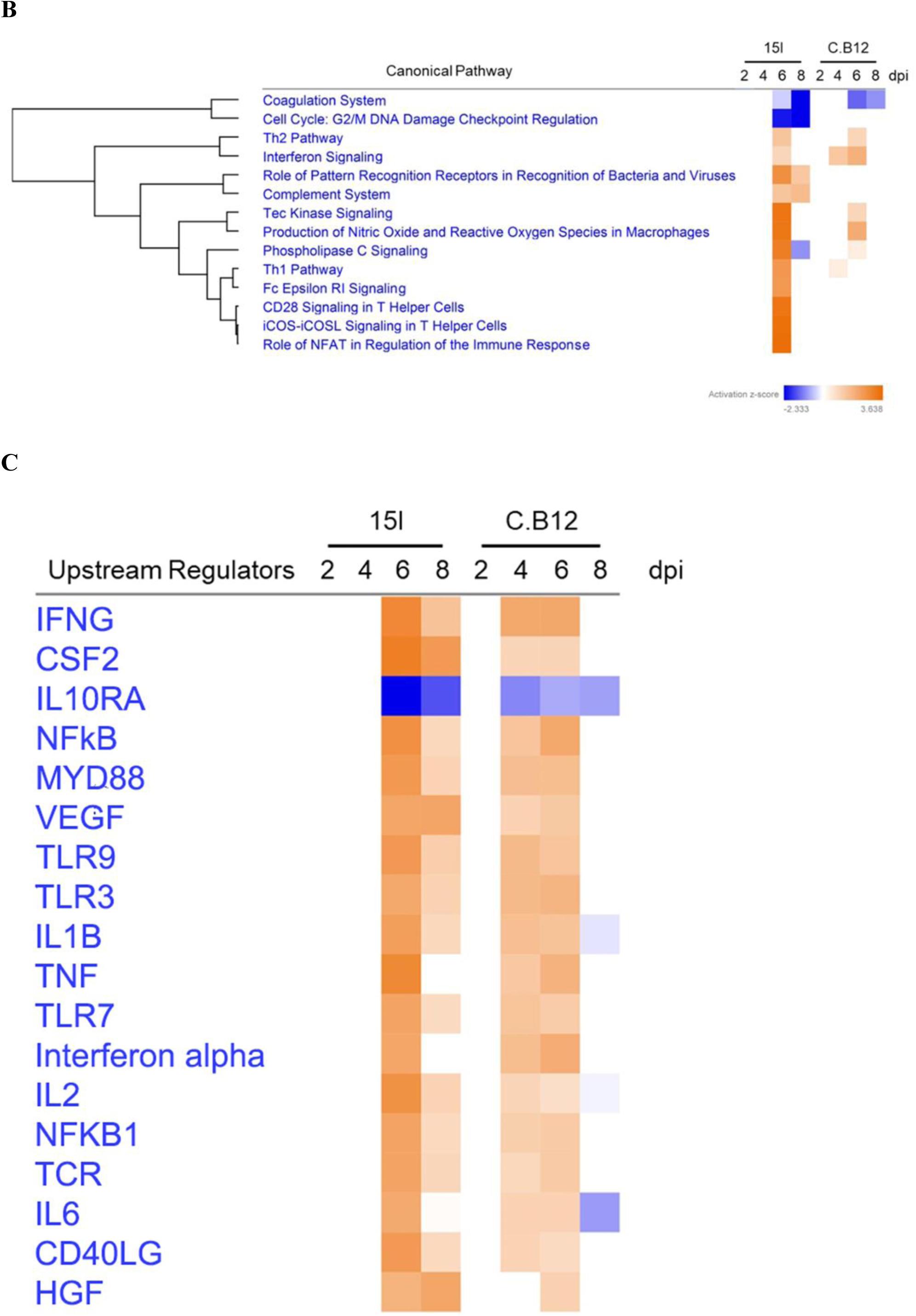
Comparison of line 15I and C.B12 chickens during *E. maxima* infection. Heatmap (A) showing genes that presented the highest mean fold difference between lines. Functional pathway analysis (B) and predicted upstream regulators in both lines (C) are presented with colour based on Z-score; orange indicating activated pathways or regulators and blue indicating de-activated pathways or regulators.

To further investigate the differences between the two chicken lines, we compared pathways enriched in each line using IPA software (Figure 4B). This highlighted commonalities and differences between the responses of the lines. The coagulation pathway was downregulated in both lines at 6 and 8 dpi, while genes associated with cell cycle regulation were uniquely downregulated in line 15I. The Th2 and the Tec kinase signalling pathways were upregulated in both lines at 6 dpi, as was interferon signalling, although the latter pathway was significantly enriched already at 4 dpi in line C.B12 chickens. Pathways that showed a stronger enrichment in line 15I compared to C.B12 chickens included the Th1, T helper cell and complement pathways. Analysis of the predicted upstream regulators revealed that both chicken lines share many of the same upstream regulators including *IFNG*, *CSF2* and vascular endothelial growth factor A (*VEGF*) although the activation of these generally occurred at 4 and 6 dpi in line C.B12 and at 6 and 8 dpi in line 15I chickens (Figure 4C).

A previous genome-wide association study using an F2 intercross between lines C.B12 and 15I revealed a 35 MB region of chromosome 2 is significantly associated with resistance to coccidiosis (31, 32). We identified genes in this region that were differentially regulated at one or more time points (Table S6). Forty-seven genes in this region were differentially regulated in at least one condition. Out of 47, 10 genes were differentially expressed between two lines, including ATP binding cassette subfamily A member 13 (*ABCA13H*), Dermatan sulphate epimerase like (*DSEL*), Serpin family B member 2 (*SERPINB2*) and Sad1 and UNC84 domain containing 3 (*SUN3*). The F-box protein 15 (*FBXO15*), which is involved in the MHC class I processing pathway, was downregulated earlier in the line C.B12 compared to line 15I chickens during *E. maxima* infection. The interferon alpha inducible protein 6 (*IFI6*), which plays a role in cell apoptosis, was upregulated in line C.B12 at 6 dpi but not in line 15I chickens, compared to non-infected chickens.

### Differential kinetics of IFN-γ and IL-10 expression in the jejunum of relatively resistant and susceptible chickens following E. maxima infection

During the analysis of RNA-Seq results, we noticed that *IFNG* and *IL10* expression increased more rapidly post-infection in line C.B12 (4 dpi) compared to line 15I (6 dpi) (Figure 5A). Expression of *IFNG* was significantly increased (FDR < 0.05) at 6 and 8 dpi in line 15I, but in line C.B12 at 4, 6 and 8 dpi, while *IL10* was significantly upregulated at 6 and 8 dpi in line 15I and at 6 dpi in line C.B12. Although *IL10* was not significantly upregulated at 4 dpi in line C.B12, this is likely due to the high variance between birds at this time point, with some samples showing elevated *IL10* counts.

**Figure 5:**
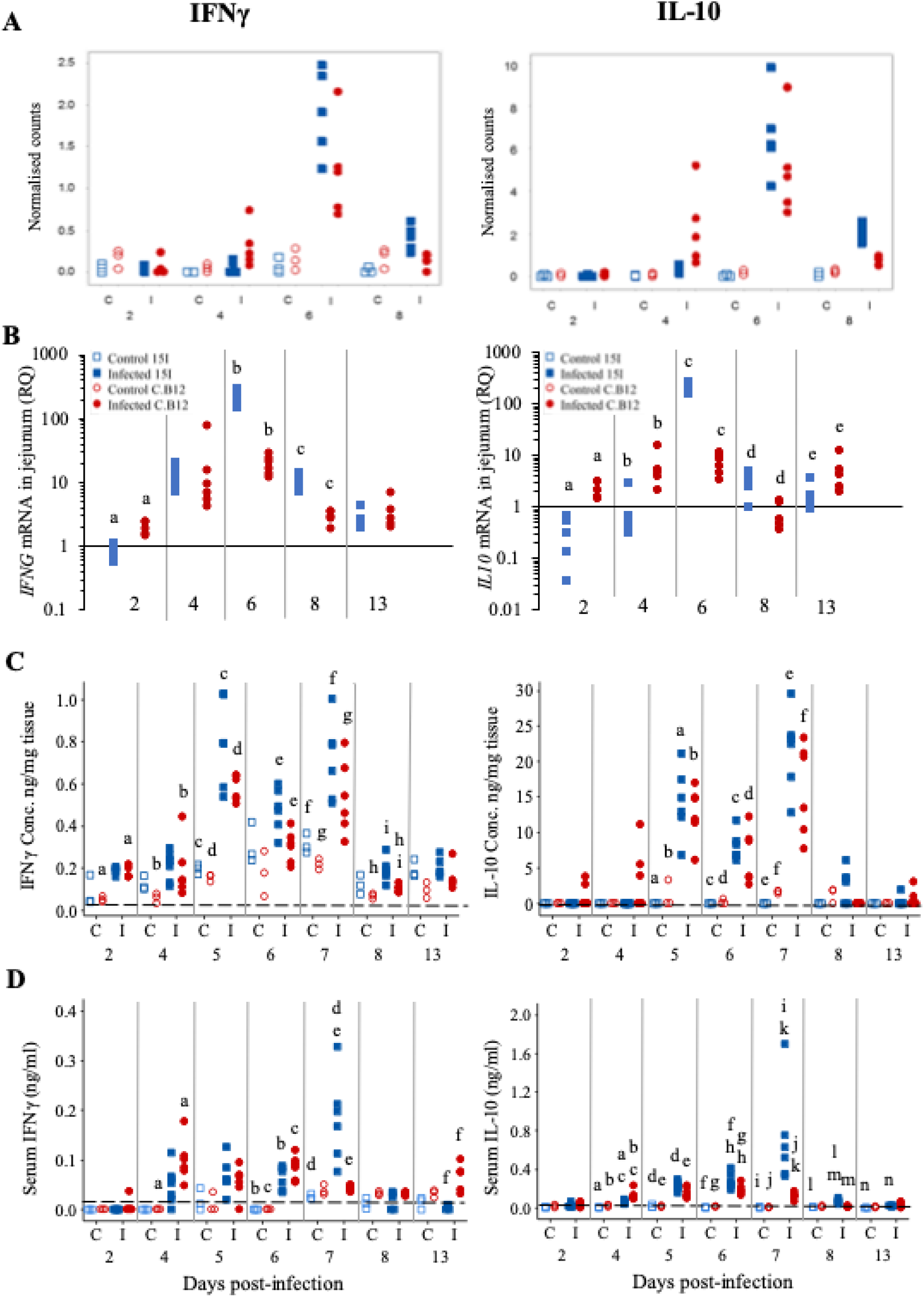
Kinetics of *IFNG* and *IL10* mRNA transcription by RNA-Seq (A) and RT-qPCR (B), protein levels in the jejunum (C) and protein levels in the serum (D) of *E. maxima*-infected chickens. Three-week old birds were orally inoculated with 100 oocysts of *E. maxima* (solid markers) or sterile water (control birds; hollow markers) and jejunum and serum samples collected at various days post-infection as indicated. Data are presented as individual birds. For RT-qPCR data, the relative quantity (RQ) of mRNA transcription of individual infected birds was calculated relative to the mean of control birds of the same line at individual time points and normalised using the 28S reference gene. Matching letters denote significant differences between groups on the same day (*p*<0.05, *n*=3 for control and *n*=5 for infected groups). C; control. I; infected. Line C.B12 shown as red circles, line 15I as blue squares.

To verify the transcriptomic results and to obtain insight into the role of IFN-γ and IL-10 in susceptibility to *E. maxima* infection, *IFNG* and *IL10* mRNA levels in lines C.B12 and 15I were determined in the jejunum at 2, 4, 5, 6, 7, 8 and 13 dpi (Figure 5B). Across control birds of all time points, line C.B12 birds had significantly higher (*p*<0.01) *IFNG* mRNA transcription in the jejunum compared to line 15I. In both chicken lines, the greatest increase in *IFNG* mRNA transcription, relative to control birds of the same line, was at 6 dpi (Figure 5B). At 6 and 8 dpi, line 15I exhibited significantly greater increases in *IFNG* mRNA levels compared to line C.B12 chickens. Analysis of IFN-γ protein in the jejunum by ELISA revealed biphasic increases in IFN-γ production at 5 and 7 dpi in both chicken lines (Figure 5C). During *E. maxima* infection, line C.B12 exhibited significantly increased IFN-γ protein in the jejunum at 2, 4, 5, 7 and 8 dpi, whereas line 15I had significantly increased IFN-γ protein at 5 and 7 dpi compared to their non-infected counterparts. Following infection, line 15I birds exhibited higher levels of IFN-γ protein in the jejunum at 6 and 8 dpi compared to line C.B12.

At 4 dpi, line C.B12 transcribed higher levels of *IL10* mRNA in the jejunum relative to age-matched control birds of the same line and control or infected line 15I birds (Figure 5B). However, the transcription of *IL10* mRNA in the jejunum of line 15I was dramatically increased, relative to controls, at 6 dpi, whereas line C.B12 expressed similar increases in *IL10* mRNA levels at 2, 4, 6 and 13 dpi. There was no significant difference in the basal transcription of *IL10* between control birds of the two lines across all time points. Similarly to IFN-γ protein levels in the jejunum, there were two peaks of IL-10 protein levels in the jejunum at 5 and 7 dpi (Figure 5C). The levels of IL-10 protein in the jejunum of the control birds was either lower than the limit of detection (as in line 15I) or very little was present (as in line C.B12) across all time points. At 5 dpi, there were significantly increased IL-10 protein levels in the jejunum of both lines of chicken. The increased IL-10 protein induced by *E. maxima* infection then slightly decreased at 6 dpi, but increased again at 7 dpi. Unlike mRNA levels, there was no significant difference in IL-10 protein levels between the two lines at any of the time points.

We also measured mRNA levels of Th17-associated genes *IL17A*, *IL17F*, *IL21*, *IL2* and *IL6* (Figure 6 and Figure S1). Although the expression of *IL2* mRNA levels at 2 and 6 dpi seemed to be upregulated, the change was not significant due to the high variance between chickens (Figure S1). Among the measured genes, only *IL21* showed a significant increase in the jejunum of both lines of chickens during *E. maxima* infection (at 4 and 6 dpi) compared to their non-infected counterparts (Figure 6). Additionally, at 4 and 6 dpi, line 15I transcribed significantly higher *IL21* mRNA levels compared to line C.B12 chickens whereas at 13 dpi, *IL21* mRNA was higher in line C.B12 birds. However, the protein levels of IL-21 in jejunum and serum were either very low or below the detection limit of the ELISA.

**Figure 6.**
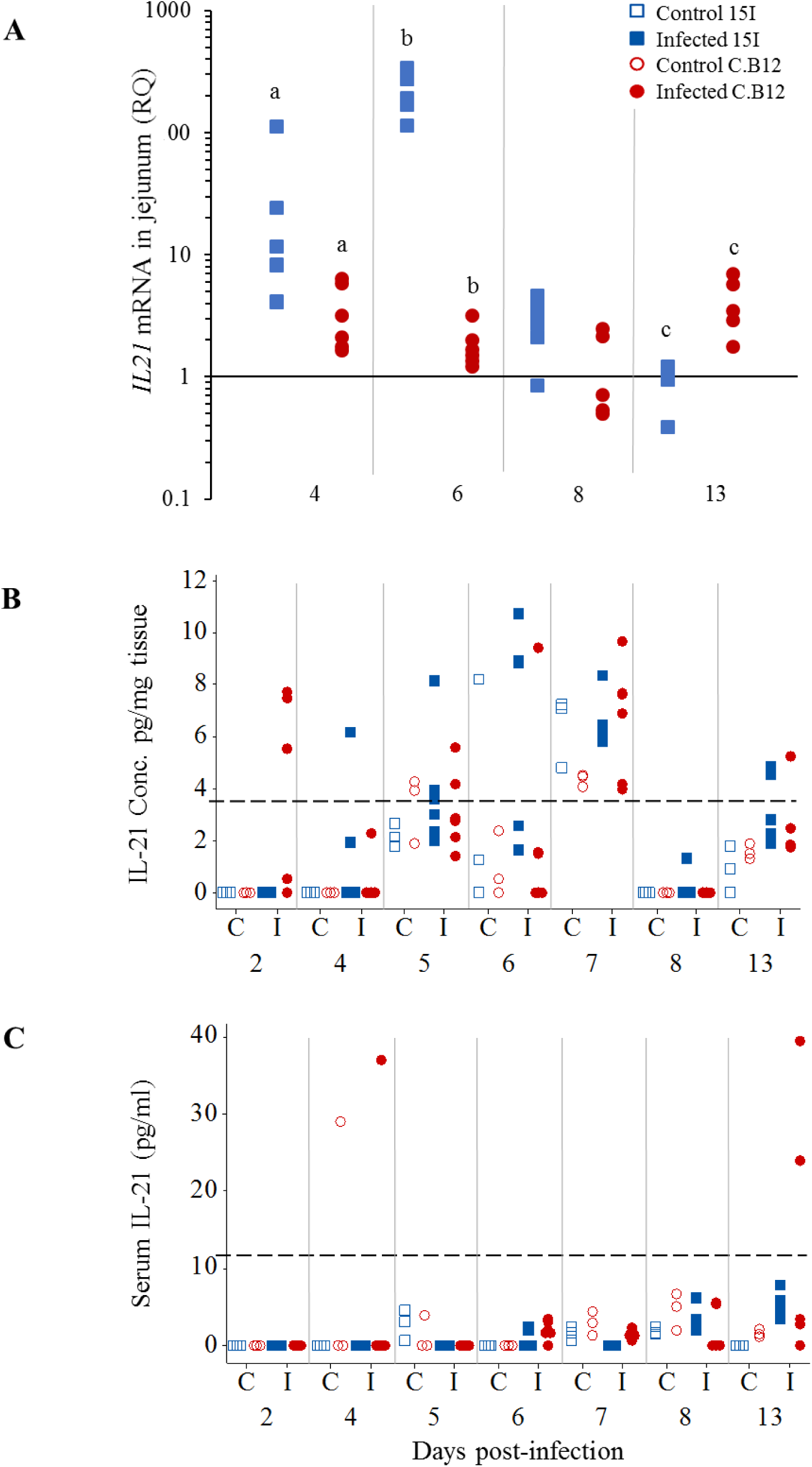
Kinetics of *IL21* mRNA transcription (A) and protein expression (B) in the jejunum and IL-21 protein levels in the serum (C) of *E. maxima*-infected chickens. Three-week old birds were orally inoculated with 100 oocysts of *E. maxima* or sterile water (control birds) and jejunum and serum samples collected at various days post-infection as indicated. Data are presented as individual birds. For RT-qPCR data, the relative quantity (RQ) of mRNA transcription of individual infected birds was calculated relative to the mean of control birds of the same line at individual time points and normalised using the 28S reference gene. Matching letters denote significant differences between groups on the same day (*p*<0.05, *n*=3 for control and *n*=5 for infected groups). C; control. I; infected.

### Differential kinetics of IFN-γ and IL-10 levels in the serum of relatively resistant and susceptible chickens following E. maxima infection

Unlike the levels of IFN-γ protein in the jejunum, the kinetics of serum IFN-γ differed between the lines with C.B12 peaking at 4 and 6 dpi and line 15I at 7 dpi (Figure 5D). *E. maxima*-infected line C.B12 exhibited significantly higher levels of serum IFN-γ at 4, 6 and 13 dpi compared to non-infected chickens. In line 15I, significantly higher serum IFN-γ was observed at 6 and 7 dpi compared to non-infected chickens. Compared to infected line C.B12, infected line 15I chickens had significantly higher serum IFN-γ at 7 dpi.

Serum IL-10 levels were significantly increased in line C.B12 following *E. maxima* infection at 4, 5, 6 and 7 dpi, while in line 15I, significantly increased serum IL-10 was observed from 4 to 13 dpi during *E. maxima* infection (Figure 5D). At 4 dpi, line C.B12 had higher levels of serum IL-10 compared to line 15I following *E. maxima* infection. However, serum IL-10 levels in the *E. maxima*-infected line 15I were significantly higher than that of C.B12 chickens at 6, 7 and 8 dpi. The levels of serum IL-10 in line 15I were dramatically increased at 7 dpi during *E. maxima* infection, whereas in line C.B12 chickens, serum IL-10 levels remained similar to those observed at 5 and 6 dpi.

### Correlation between local and systemic IFN-γ and IL-10 production and parasite burden

To investigate the effect of IFN-γ and IL-10 on *E. maxima* burden, the correlation between jejunum and serum IFN-γ and IL-10 protein levels and *E. maxima* replication were calculated (Table 2). Both local (jejunum) and systemic (serum) IFN-γ and IL-10 levels in both lines of chickens correlated positively with *E. maxima* burden. Serum IFN-γ in line 15I correlated more strongly with *E. maxima* burden than in line C.B12 chickens, whereas tissue IFN-γ correlated more strongly with *E. maxima* burden in line C.B12 compared to line 15I chickens. Both serum and jejunum IL-10 in line 15I correlated more strongly with *E. maxima* burden compared to line C.B12 chickens. We also measured the effect of IFN-γ and IL-10 on BWG. Although the expression of IFN-γ and IL-10 in the jejunum and serum correlated negatively with BWG, the correlation was not significant (*p* > 0.05) (data not shown).

**Table 2.** Correlation of IFN-γ and IL-10 in serum and jejunum with E. maxima replication.

|  |  | Line 15I | Line C.B12 |
| --- | --- | --- | --- |
| IFN-γ | Serum | 0.52<br>(p<0.001) | 0.19<br>(p=0.25) |
|  | Tissue | 0.51<br>(p<0.001) | 0.63<br>(p<0.001) |
| IL-10 | Serum | 0.74<br>(p<0.001) | 0.61<br>(p<0.001) |
|  | Tissue | 0.64<br>(p<0.001) | 0.57<br>(p<0.001) |

### Cellular changes following E. maxima infection

To investigate and compare changes to the immune cell populations in the two lines of chickens at the early stages of *E. maxima* infection, IHC was performed with jejunum collected at 4 dpi (Figure 7). We first compared jejunum of uninfected birds to establish if intrinsic differences between the lines existed. There was no significant difference in the number of cells expressing any of the measured cell markers in the villus lamina propria (Figure 7A) or epithelium (Figure 7B) between the two lines, although line C.B12 displayed slightly higher numbers of CD4^+^, CD8α^+^, γδ T^+^ and αβ1^+^ T cells.

**Figure 7.**
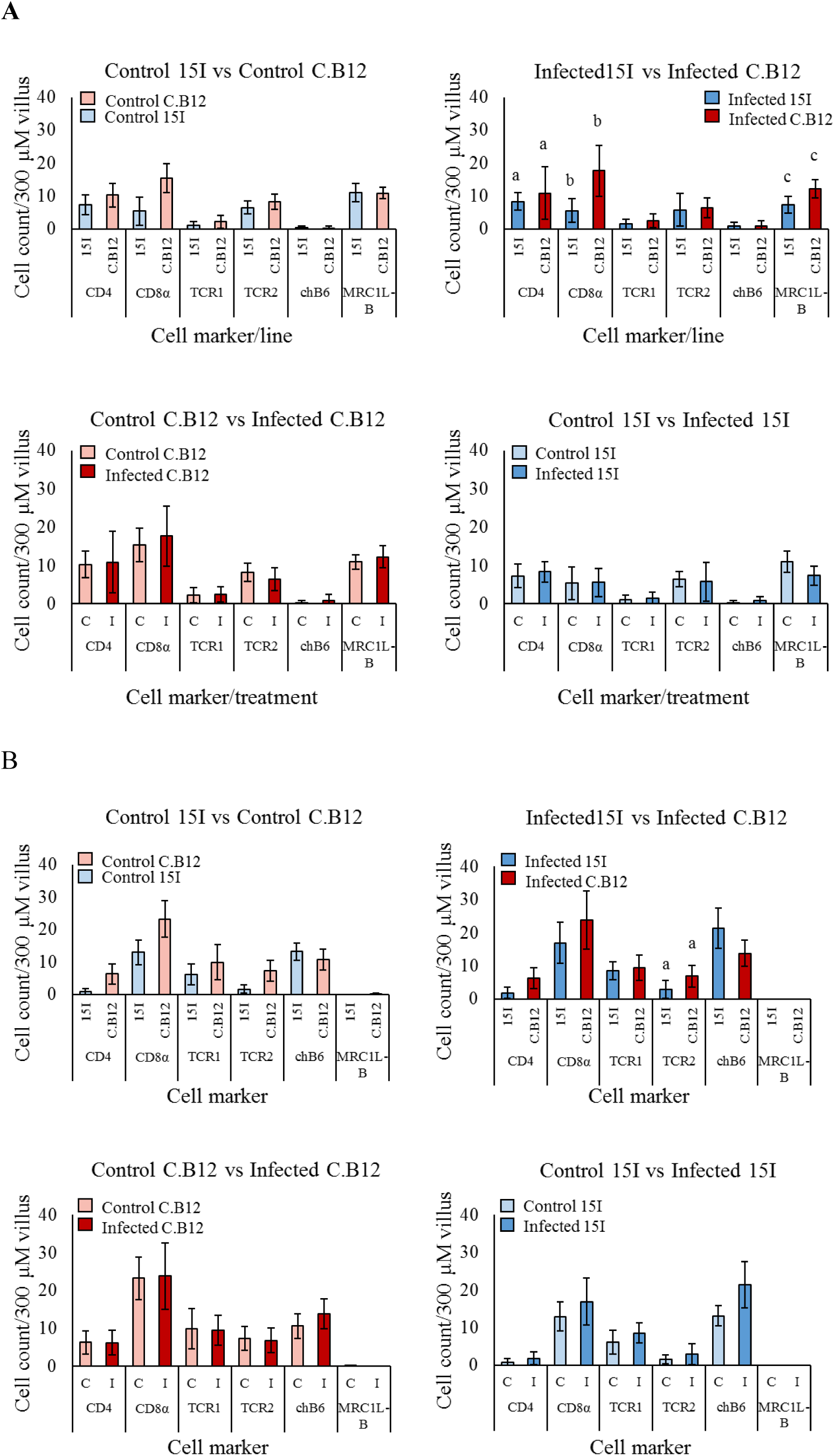
Populations of CD4, CD8α, γδ T cells, αβ1 T cells, chB6 and MRC1L-B LPLs (A) and IEL (B) in the jejunal villi of line C.B12 and line 15I chickens at 4 dpi with *E. maxima*. Shown are data comparing control birds of both lines, infected birds of both lines, control and infected line C.B12 birds and control and infected line 15I birds. LPLs and IEL were counted from nine villi of one section per bird (*n*=3 for uninfected and *n*=5 for infected groups). Each bar represents the mean number of cells per 300 μm of villus (± SD). Matching letters denote significant differences between groups (*p*<0.05). C; Control. I; infected.

At 4 dpi, there was no difference in the population of the measured cell markers between *E. maxima*-infected and non-infected chickens in the jejunum lamina propria or epithelium of either chicken line. However, comparison of the number of cells in *E. maxima*-infected tissues revealed significantly lower numbers of CD4^+^, CD8α^+^ and MRC1L-B^+^ cells in the lamina propria (Figure 7A) and αβ1^+^ T cells in the epithelium of the villi (Figure 7B) in line 15I compared to line C.B12 chickens.

We also measured changes to the immune cell populations in the lamina propria and epithelium of the crypts in both lines of chickens (Figure S2). There was no significant difference in the number of cells between uninfected chickens of line C.B12 and line 15I, or between *E. maxima*-infected and non-infected chickens within each lines. Comparison of the number of cells in the crypts of *E. maxima*-infected chickens revealed significantly lower numbers of αβ1^+^ T cells and higher numbers of chB6+ cells in the epithelium of line 15I compared to line C.B12 chickens (Figure S2B).

## Discussion

Understanding the basis of resistance to *E. maxima* is important for the commercial poultry industry as it would enable identification of quantifiable resistance or susceptible phenotypes, allowing for the selective breeding of chickens for resistance against this and possibly other *Eimeria* species. Thus, investigation of host responses to *Eimeria* infection in the relatively resistant and susceptible White Leghorn chicken lines C.B12 and 15I has important economic implications for the poultry production industry, in addition to avian well-being and food security. In this study, we characterised the kinetics of differential gene expression in these two lines of chicken, as well as the kinetics of local and systemic protein expression and mRNA transcription of IFN-γ, IL-10, IL-21 and Th17 responses. We have also investigated cellular differences between control and infected birds of both lines during the early stages of infection. The results indicate the importance of early activation of interferon signalling pathways, with IFN-γ, IL-10 and IL-21 responses during the innate phase of infection associated with resistance to *E. maxima*. This research builds on previous work, investigating the importance of these responses from transcriptome to protein levels in the jejunum, the site of *E. maxima* infection, and systemically at the protein level in the serum.

Transcriptomic analysis of jejunal tissue from chicken lines C.B12 and 15I infected with *E. maxima* revealed differences in the kinetics of the host immune response and provided information on the different biological pathways involved. Commonalities between the two lines included strong upregulation of *IFNG,* various chemokines and complement components at 6 dpi, which agrees with previous transcriptome based analysis of chicken caecal epithelial responses to *E. tenella* (33). Although there was no difference in *E. maxima* replication at 4 dpi between the two lines, early immune responses observed in relatively resistant line C.B12 at this time point, in particular interferon responses, may be sufficient to reduce *E. maxima* replication at 7 dpi compared to line 15I where these responses did not occur until 6 dpi, potentially leading to a delay in the inhibition of *E. maxima* replication. Pathways involved in Th1 and Th2 responses were also upregulated at 4 dpi in line C.B12. Although 4 dpi is likely too early for such adaptive responses, higher numbers of CD4, CD8α and αβ1 T cells were present in the jejunum of control and infected line C.B12 compared to line 15I birds, and are cell types associated with these responses. Regardless of resistance and susceptibility to *E. maxima*, both chicken lines share many of the same upstream regulators including IFN-γ, IL-10RA and IL-2 that may cause changes in gene expression; however, similar to functional pathway analysis, all the predicted upstream regulators affect expression in line C.B12 at 4 dpi and 6 dpi, whereas line 15I chickens are not affected by the same upstream regulators until 6 dpi, supporting the importance of the early immune responses in resistance to *E. maxima* infection. Transcriptomic analysis also revealed a set of interferon-stimulated genes that were uniquely responding in line C.B12, including *MX1*, *RSAD2* and *OASL*, that may be involved in the relative resistance displayed by line C.B12.

One of the important findings of this study was that higher increases in early (2 and 4 dpi) IFN-γ and IL-10 production correlated with resistance to *E. maxima*, whereas a more gradual increase (a minor increase at 2 and 4 dpi but a dramatic increase at 6 and 8 dpi) in production of these cytokines was correlated with susceptibility, indicating the timing at which the immune response is mounted is paramount to resistance. These results were evaluated by IHC, showing that an intrinsically higher presence of IFN-γ-producing (CD4^+^, αβ1^+^ T cells and MRC1L-B^+^ macrophages (34)) LPL and IEL were present in relatively resistant line C.B12 in the villi of control birds than in line 15I at 4 dpi. Moreover, significantly higher numbers of MRC1L-B+ macrophages, CD4+ and CD8α+ cells were detected in the lamina propria of infected line C.B12 compared to line 15I birds, indicating macrophage and NK cell involvement at 4 dpi. Chicken intestinal IEL include NK cells which may express CD8α (35), chB6 (36) or TCRγδ (37). Likewise, Wakelin et al. (39) showed Con A-responsive cells in the mesenteric lymph nodes appeared earlier and produced more IFN-γ in *E. vermiformis*-resistant mice following infection. Taken together, significantly up-regulated IFN-γ expression in the jejunum of *E. maxima*-infected chickens is likely due to the recruitment and stimulation of MCR1L-B^+^, CD4^+^ and CD8α^+^ cells. Hong et al., (19) showed that *IL10* and *IFNG* mRNA transcription was robustly increased at 4 and 6 dpi in CD4 and CD8 cell subpopulations following *E. maxima* infection. The current study identified higher numbers of CD4 and CD8α IEL and LPL in line C.B12, both prior to and following infection. In support of these findings, higher numbers of CD4^+^ IEL were detected in the duodenum during early *E. acervulina* infection in resistant chickens (40) and increased CD4^+^ LPL were detected within 24 h of intra-caecal inoculation of *E. tenella* sporozoites (41), implying CD4^+^ cells are effectors of *Eimeria* resistance early on during infection and could be a source of the early IL-10 and IFN-γ observed in this study.

IL-10 is a pleiotropic cytokine and in addition to maintaining the Th1/Th2 balance, it is also important to normal gut homeostasis, regulating NK cell and macrophage activity, limiting proinflammatory cytokine production and promoting epithelial cell proliferation amongst other functions (42). The impact of IL-10 on the outcome of *Eimeria* infection is likely dependent on both the timing and magnitude of its production. Early IL-10 may be involved in mediating innate responses; pegylated recombinant human IL-10 induces IFN-γ, perforin and granzyme B secretion in CD8^+^ T cells (43). Other publications have indicated that IL-10 reduces the efficacy of the immune response to *Eimeria*. Antibody-mediated IL-10 depletion in broilers enhanced weight gain and decreased oocyst production following inoculation with an attenuated *Eimeria* spp. vaccine (*E. maxima*, *E. tenella* and *E. acervulina)* (30) and did not appear to affect adaptive immunity as IL-10-depleted-chickens displayed similar weight gains following vaccination then challenge as control birds (44). Additionally, in broilers treated with CitriStim, a yeast mannan-based feed additive, and given an attenuated vaccine (*E. maxima*, *E. tenella* and *E. acervulina*), reduced *IL10* mRNA was found in the caecal tonsils which was accompanied by reduced oocyst shedding and improved feed efficiency and weight gains (45). Although in the study by Rothwell et al. (27), *IL10* transcripts were detected in the spleen of control birds, we did not detect IL-10 at a protein level in the serum in our study. Rothwell et al. (27) also observed extremely low basal levels of *IL10* mRNA in the jejunum of uninfected chickens whereas the current results suggest that *IL10* mRNA is transcribed in the jejunum under normal homeostatic conditions. This discrepancy is attributable to the increased sensitivity of the primer and probe sequences used in this study (data not shown). Levels of IL-10 protein positively correlated with *E. maxima* replication and it is plausible *E. maxima* is inducing IL-10 as an immune evasion strategy. Similar to the findings by Hong et al. (19), each vaccination with *E. maxima* led to increased serum IL-10, however the extent to which increases were observed gradually decreased with each subsequent vaccination, whereas serum IFN-γ was only increased after the first vaccination (in our unpublished data). The current study suggests that an early, modest induction of IL-10 does not negatively impact resistance to *E. maxima* infection, but excessive IL-10 production disrupts the efficacy of the protective response. These findings imply that IL-10 can be suitable as a biomarker of susceptibility at late time points with *E. maxima* infection, but less suitable as a predictor of susceptibility prior to or early on during infection.

IL-17A and IL-17F are mainly considered cytokines of the Th17 cell lineage, which functions in autoimmune disease and defence against bacterial, fungal and parasitic pathogens (46, 47, 48). More recently IL-17A and IL-17F have been related to innate cells including NK and γδ T cells and macrophages. They are important mediators of mucosal immunity and innate responses, with functions including neutrophil recruitment, macrophage activation and IFN-γ production and chemokine and antimicrobial peptide production in epithelial cells (49, 50). As our study implies, early innate responses are key to resistance to *E. maxima* and previous studies have indicated that *Eimeria* spp. infection in chickens leads to the increased transcription of *IL17A*, as well as *IL2* and *IL6* mRNAs (19, 51). In contrast, our RT-qPCR and RNA-Seq data revealed there was no significant change in *IL17A*, *IL17F*, *IL2* and *IL6* mRNA levels during *E. maxima* infection. Although *IL17A* and *IL2* mRNA levels at 2 and 6 dpi seemed to be upregulated, the change was not significant due to the high variance between chickens within the same group. Previously it has been suggested that IL-17A impairs immunity to *Eimeria* spp. infection. Zhang et al. (52) showed increased *IL17A* mRNA transcription at 6 hours post infection with *E. tenella*. IL-17A depletion reduced heterophil infiltration and associated immunopathology in the caeca, but also reduced oocyst output indicating that IL-17A is involved in susceptibility to *E, tenella.* In addition, Del Cacho et al. (53) also found that IL-17A reduced *E. tenella* schizont development and migration. Among the Th17-associated genes tested, only *IL21* mRNA levels were increased in the jejunum of both lines of chickens during *E. maxima* infection compared to non-infected chickens. A member of the IL-2 family, IL-21 plays important roles not only in Th17 differentiation, but also in innate immunity, with functions including enhancement of cytoxicity and IFN-γ production in NK and CD8 T cells (54, 55). Additionally, IL-21 plays key roles in autoimmune disease and in shaping humoral and cellular immune responses to parasitic infection (56, 57). In chickens, increased *IL21* mRNA levels are reported in autoimmune vitiligo. To date, IL-21 has not been previously found to have a role during *Eimeria* infection. The kinetics of our study revealed that the pattern of *IL21* mRNA transcription was similar to *IFNG* and *IL10* in the jejunum, indicating that IL-21 may also be involved in resistance to *E. maxima* through mediating innate immunity. Similar transcription patterns of *IFNG*, *IL10* and *IL21* mRNA were reported during the development of vitiligo lesions (58). Moreover, in mice IL-21 modulates differentiation of CD4 and CD8 T cell subsets in a context-dependent manner and certain cytokines, including IL-10, may compensate for IL-21 (59). Since *E. maxima* infection leads to an increase in CD8α T cell numbers, it is possible that the co-expression of IL-21, IFN-γ and IL-10 may play an important role in the enhancement of CD8 T cell responses, as reflected in the higher numbers of CD8α IEL and LPL in the jejunum of line C.B12 birds observed in this study. Previously, cytotoxic CD8 cell activity was shown to be a component of protective immunity to secondary *E. tenella* infection (41, 60, 61) and resistance and IFN-γ production during primary *E. acervulina* infection in chickens (62). The early timing of this response in line C.B12, alongside the fact that no other Th17-associated genes tested were changed during infection, indicates that Th17 responses are not involved during *Eimeria* spp. infection.

The present study suggests that the timing of the immune response is crucial for *E. maxima* resistance. Immunity to *Eimeria* arises during sporozoite translocation through the lamina propria in chickens (60, 63, 64). Therefore logically, resistance to *Eimeria* spp. relies on the host response in the first few days of infection, when the majority of sporozoites are present in the lamina propria and in contact with LPL. The increased IL-10 observed in line 15I in the serum suggests that systemic IL-10 production promotes susceptibility to *E. maxima*, but given the positive correlation of IL-10, IFN-γ and IL-21 with one another and the higher expression in resistant chickens at early time points implies that the balance between the three is imperative for effective immunity to *E. maxima*.

## Methods

### Ethics statement

Animal work was carried out in strict accordance with the Animals (Scientific Procedures) Act 1986, an Act of Parliament of the United Kingdom, following approval by the Royal Veterinary College Ethical Review Committee and the United Kingdom Government Home Office.

### Animals and parasites

Chickens of two inbred White Leghorn lines were used in this study. Inbred line 15I, relatively susceptible to *E. maxima* infection, originate from the Regional Poultry Research Laboratory (East Lansing, MI). Reaseheath C (line C, C.B12) chickens, relatively resistant to *E. maxima* infection, originate from the University of Cambridge (Cambridge, UK). Both flocks were maintained at the National Avian Research Facility (NARF; The Roslin Institute, UK).

The Weybridge (W) strain of *E. maxima* was used (65). Parasites were passaged at frequent intervals through dosing and faecal recovery as described previously (66), and used less than one month after sporulation.

### Experimental design, sampling and data collection

Line C.B12 and 15I chickens were supplied at day-of-hatch without prior vaccination to the Royal Veterinary College, where chickens were reared in coccidia-free, environmentally enriched conditions with feed and water provided *ad libitum*. Chickens were housed following Defra stocking density guidelines and raised under industry-standard conditions. Prior to inoculation, chickens (*n* = 60 and *n* = 62 for lines C.B12 and 15I, respectively) were randomly allocated to four different pens corresponding to the two lines and two different experimental treatments: control and infected. The absence of prior coccidian infection was confirmed by faecal flotation. Three-week-old chickens were orally infected with 100 sporulated *E. maxima* oocysts (test) or sterile water (control).

In order to analyse differential kinetic immune responses elicited by *E. maxima* infection, blood and small intestine (jejunum) were collected from 3 chickens in the control groups and a minimum of 5 chickens in the infected groups at 2, 4, 5, 6, 7, 8 and 13 dpi. Body weight was recorded individually two days prior to infection and prior to culling at each sampling point and the percentage weight gain calculated. Chickens were culled by cervical dislocation following the Schedule 1 method, and death confirmed by permanent cessation of circulation. Blood was collected from the jugular vein immediately after culling. For serum, blood samples were allowed to clot at room temperature, followed by centrifugation at 1,500 x *g* for 3 min and the separated serum stored at -20°C.

Approximately 10 cm of small intestine, spanning 5 cm anterior and posterior to Meckel’s diverticulum (the mid-point of the intestinal area infected by *E. maxima*) was excised (66), and parasite-related lesions scored as described by Johnson and Reid (67). Approximately 0.5 cm of jejunum, 1 cm anterior to the Meckel’s diverticulum, was collected into RNA*later*® Stabilization solution (Life Technologies, CA, USA) for gene expression analysis and by snap-freezing in liquid nitrogen for analysis of tissue protein levels. For histology, 1 cm of jejunum tissue was snap frozen in optimum cutting temperature (OCT) compound on liquid nitrogen and stored at -80°C until use. For parasite quantification, the remaining excised tissues were stored in RNA*later*® Stabilization solution at 4°C overnight then at -20°C after removal of the reagent.

### Isolation of genomic DNA and quantitative PCR (qPCR) for E. maxima replication

Total genomic DNA (gDNA) was isolated from the excised small intestine as described previously (68). Briefly, tissue samples were weighed and suspended in an equal volume (w/v) of tissue lysis buffer (Buffer ATL, Qiagen, Crawley, UK), and homogenized employing a TissueRuptor (Qiagen). Subsequently, the equivalent of ≤ 25 mg of the homogenate was used to carry out the gDNA isolation using a DNeasy® Blood and Tissue kit (Qiagen) according to the manufacturer’s instructions. The gDNA was stored at -20°C, until further investigation.

Quantitative PCR (qPCR) was performed as previously described (68) using a CFX96 Touch® Real-Time PCR Detection System (Bio-Rad Laboratories, CA, USA). For the quantification of *E. maxima* total genome copy numbers, we used the primers EmMIC1_For (forward: 5’-TCG TTG CAT TCG ACA GAT TC-3’) and EmMIC1_Rev (reverse: 5’-TAG CGA CTG CTC AAG GGT TT-3’) (10). The chicken cytoplasmic β-actin (actb) gene was used for data normalization, amplified using the primers actb_FW (forward: 5’-GAG AAA TTG TGC GTG ACA TCA-3’) and actb_RV (reverse: 5’-CCT GAA CCT CTC ATT GCC A-3’) (69). Briefly, each sample was amplified in triplicate in a 20 µL volume containing 1 µL of total gDNA, 300 nM of each primer, 10 µL of SsoFast™ EvaGreen® Supermix (Bio-Rad Laboratories), and 8.8 µL of nuclease-free water (Life Technologies) with qPCR cycling conditions that consisted of 95°C for 2 min as initial denaturation, followed by 40 cycles of denaturation at 95°C for 15 sec and annealing/extension at 60°C for 30 sec. Dissociation curves were generated to analyse individual PCR products after 40 cycles. Each qPCR assay included the relevant gDNA dilution series as standards (68) and no template controls. The genome copy numbers from the chicken (actb) and the *E. maxima* parasites (EmMIC1) were estimated by comparison with the gDNA dilution series. Triplicate data arising from each test sample were averaged and standardized by comparison with the concentration of chicken genome as a ratio of *E. maxima* genomes/chicken genomes.

### Total RNA preparation and quantitative real-time PCR (RT-qPCR)

Total RNA was extracted from the jejunum using the RNeasy® Mini Spin Column Kit (Qiagen) following the manufacturer’s instruction. Briefly, approximately 25 mg of tissues were homogenized in 2 mL tubes containing 600 μL of Buffer RLT with 2% β-mercaptoethanol and a stainless steel bead (5 mm, Qiagen) using a TissueLyser II system (Qiagen). The supernatant was collected and applied to a QIAshredder column (Qiagen) to improve the quality of total RNA. The flow-through was mixed with an equal volume of 70% ethanol and applied to an RNeasy® Spin column (Qiagen). Contaminating gDNA was digested by on-column DNase treatment using RNase-free DNase (Qiagen) and total RNA was eluted with 80 μL of nuclease-free water (Qiagen). The absorbance at 230, 260 and 280 nm was measured using a NanoDrop 1000 spectrophotometer (Thermo Scientific). For the transcriptomic study, the quantity and quality of total RNA was assessed using a Qubit® RNA BR assay kit (Life Technologies) by Qubit® 3.0 fluorometer (Life Technologies) and an RNA ScreenTape (Agilent Technologies, USA) by 2200 TapeStation System (Agilent Technologies), respectively.

The mRNA levels of target cytokines were quantified by TaqMan® real-time quantitative PCR (RT-qPCR) as described previously (70) (Table 3). TaqMan assays were performed using the One-Step RT-PCR Master Mix reagent, and amplification and detection were performed using the TaqMan Fast Universal PCR Master mix in the AB 7500 FAST Real-Time PCR System (Applied Biosystems). Standard curves for each target gene were generated as previously described (71). Each RT-qPCR assay contained triplicate no-template controls, test samples and a log10 dilution series of standard RNA. Relative gene expression of the infected birds to control birds was calculated using the Pfaffl method as described by Sutton et al. (70) and the results were presented as log10 fold-change of target gene in each line at each time point.

**Table 3.**
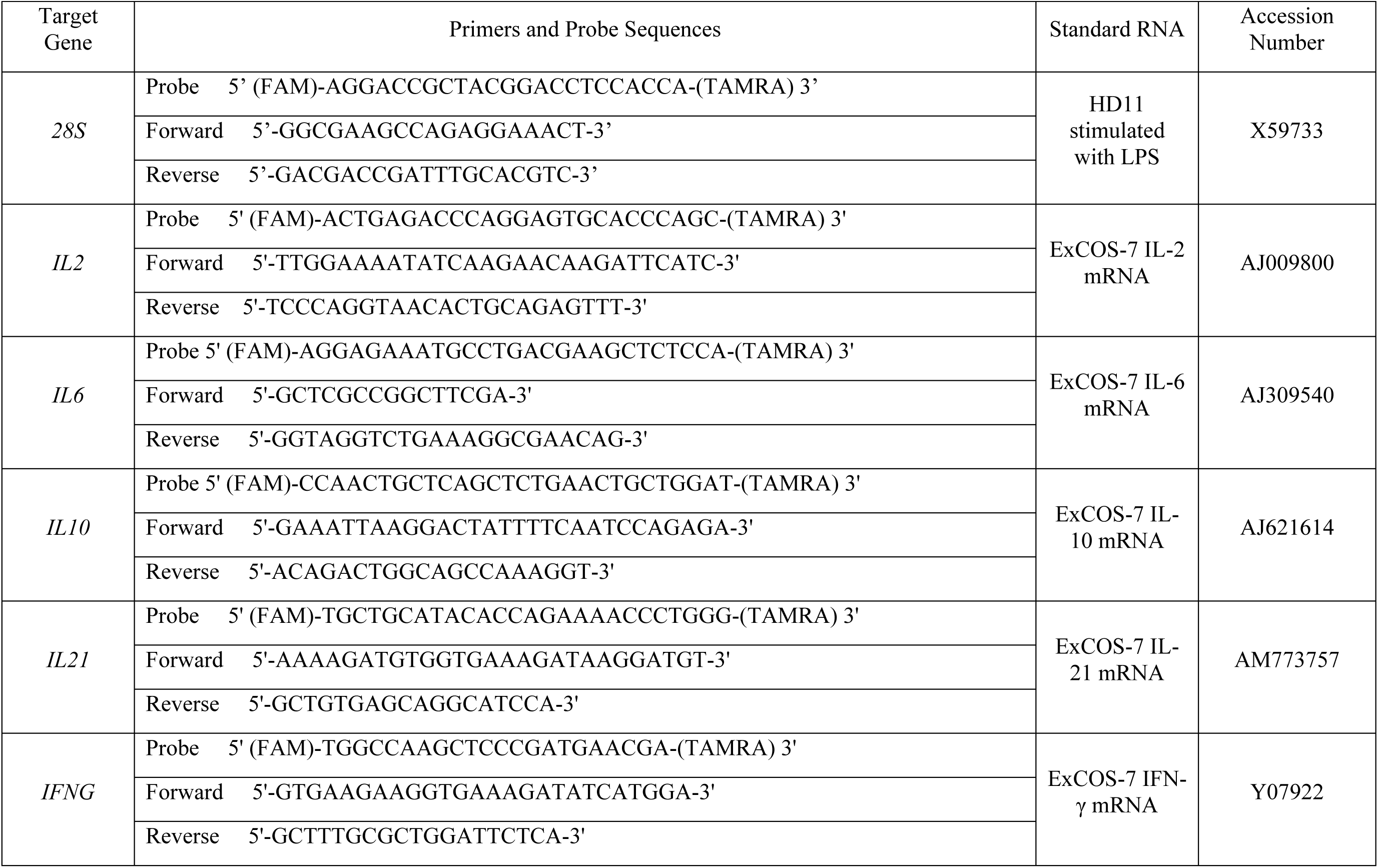
Primers and probes used in RT-qPCR.

**Table 4.** Antibodies used in IHC staining.

| Antibody | Specificity | Clone | Dilution | Reference |
| --- | --- | --- | --- | --- |
| Mouse anti-chicken CD4 | Chicken CD4 | CT-4 | 1:400 | Chan *et al.* (1988) |
| Mouse anti-chicken CD8α | Chicken CD8α | 3-298 | 1:400 | Luhtala *et al.* (1995) |
| Mouse anti-chicken TCR$\gamma\delta$ | Chicken TCR$\gamma\delta$ | TCR1 | 1:400 | Chen *et al.* (1988) |
| Mouse anti-chicken TCR$\alpha\beta$/vβ<sub>1</sub> | Chicken TCR$\alpha\beta$<sub>1</sub> | TCR2 | 1:400 | Chen *et al.* (1988) |
| Mouse anti-chicken monocyte/macrophage | Chicken mannose receptor 1 (MRC1) on monocytes, macrophages, interdigitating cells and microglia | KUL01 | 1:800 | Mast *et al.* (1998) |
| Mouse anti-chicken Bu-1/ChB6 | Chicken chB6, present on B cells and epithelial NK cells | AV20 | 1:800 | Rothwell *et al.* (1996) |

### RNA-Seq library construction, sequencing and data analysis

The total RNA of 64 samples were submitted to Edinburgh Genomics, where libraries from each of the 64 individuals were generated using automated TruSeq stranded mRNA-Seq library, and the individual jejunum transcriptomes were sequenced by 150 cycles generating paired-end reads using Illumina HiSeq 4000 technology to yield at least 290M reads. The 64 samples included 3 control and 5 *E. maxima* infected samples from lines C.B12 and 15I at 2, 4, 6 and 8 dpi.

Reads were trimmed using Trimmomatic (ver. 0.36) (72) to remove adaptor sequences of the TruSeq Stranded mRNA kit and for quality. After trimming, reads were required to have a minimum length of 75 bases. The RNA-seq reads were mapped to the reference genomes using the STAR aligner software package (ver. 2.5.1b) (73). The reference genome used for mapping was the *Gallus gallus* (Gallus_gallus-5.0) and *Eimeria maxima* (EMW001) genomes from Ensembl (https://www.ensembl.org/index.html). The annotation used for counting was derived from the *Gallus gallus* genome only, such that reads mapping to *E. maxima* were not counted in downstream analysis. Raw counts for each annotated gene were obtained using the featureCounts software (ver. 1.5.2) (74).

Differential gene expression analysis was performed using the Bioconductor edgeR package (ver. 3.16.5) (75). Statistical assessment of differential expression was carried out with the likelihood-ratio test. Differentially expressed genes were defined as those with FDR <0.05 and logFC > 1.6. Heatmaps were constructed in R using the pheatmap package. Overrepresentation of GO terms was investigated using the PANTHER Overrepresentation Test (released December 5, 2017) using Fisher’s Exact with FDR multiple test correction. Network analysis for both sample-sample and gene-gene networks was performed using BioLayout 3D (76) which performs a Pearson correlation matrix calculated for each pair of samples or genes, using a modified Fruchterman-Rheingold algorithm, with correlation cut offs of r = 0.93 (sample-sample) and r =0.87 (gene-gene). Clustering was performed on these networks using the Markov clustering algorithm (MCL) with an inflation value of 2.4 (sample-sample) and 1.4 (gene-gene). The IPA program (Ingenuity® System) was used to identify cellular canonical pathways and physiological functions that are affected by *E. maxima* infection in the host (*p*-value < 0.05 and *q*-value < 0.05).

### Preparation of protein lysates from tissue samples and capture ELISA assays

To determine protein levels of cytokines in tissues, protein lysates were prepared from the collected jejunum using the modified protein lysis buffer (20 mM Tris (pH 7.5), 100 mM NaCl, 0.5% NP-40 (IGEPAL® CA-630, Sigma), 0.5 M EDTA, 0.5 mM phenylmethylsulfonyl fluoride (Sigma) and 0.5% protease inhibitor cocktail (Sigma)). Approximately 20 mg of jejunum were mixed with 600 μL of the prepared protein lysis buffer and homogenized using 5 mm stainless steel beads (Qiagen) and a TissueLyser II system (Qiagen), twice at 25 Hz for 2 min with a 5 min incubation on ice between the two homogenizations. The samples were centrifuged at 13,000 x *g* for 10 min at 4°C and the supernatants transferred to chilled microcentrifuge tubes. The concentrations of the protein lysates were measured using the BCA Protein Assay kit (Thermo Scientific) according to the manufacturer’s instructions.

IL-10 and IFN-γ protein levels in serum and tissues were measured by ELISA. IL-10 was quantified using an in house-developed ELISA system (kindly provided by Dr. Z. Wu) for serum as described previously (29) and was adapted for use with tissue lysates. Briefly, assay plates (Nunc Immuno MaxiSorp, Thermo Scientific) were coated with 3 μg/mL of capture antibody diluted in carbonate/bicarbonate buffer at 4°C overnight. Plates were incubated with 50 μL of 2-fold serially diluted standards, sera or protein lysates for 1 hr, followed by incubation with 1 μg/mL of detection antibody for 1 hr. The plates were incubated with the Pierce High Sensitivity streptavidin-HRP (1:10,000 dilution, Thermo Scientific) for a further hour before adding 50 μL of 1-Step Turbo TMB (Thermo Scientific). After 10 min, the reaction was stopped by adding 50 μL of 2 N sulphuric acid. The absorbance was read at 450 nm (650 nm as a reference). Serum and tissue IFN-γ levels were quantified using the Chicken IFN-γ CytoSet kit (Life Technologies) as per the manufacturer’s instructions.

The standard curve was fitted to a four-parameter logistic regression curve and final concentration measures were determined using the online program provided by elisaanalysis.com (http://www.elisaanalysis.com/). The quantity of IL-10 and IFN-γ protein in the jejunum was converted from the concentration determined by ELISA to the quantity of protein in 1 mg of tissue by correcting for the amount of protein lysate used in the ELISA and the total protein lysate in 1 mg of tissue.

### Immunohistochemistry (IHC)

Immunohistochemistry was performed to determine differences in cell populations in the jejunum of line C.B12 and 15I chickens at 4 dpi with *E. maxima*. Cryostat sections (7 μm thick) were picked onto Superfrost® glass slides (Thermo Scientific) and air-dried. Sections were fixed in acetone with 0.75% H_2_O_2_ for 10 min at room temperature and air-dried for a further 5 min. The sections were incubated with monoclonal antibodies (purchased from Southern Biotech, Cambridge, UK, Table 2) specific for various leukocyte subpopulations. The Vectastain Elite ABC (Mouse IgG) Kit (Vector Laboratories, CA, USA) was used to detect monoclonal antibodies and peroxidase activity developed using the AEC staining kit (Sigma) following the manufacturer’s instructions. Subsequently, sections were counterstained with haematoxylin Z (CellPath, Newtown, UK), and bluing performed with Scott’s Tap Water (tap water, 2 % magnesium sulphate, 0.35% sodium bicarbonate). Slides were mounted in Aquamount AQ (Vector Laboratories) and images were captured with an Eclipse Ni microscope (Nikon, Tokyo, Japan), followed by quantification of the subpopulation of T lymphocytes using ZEN lite 2012 software (blue edition, Carl Zeiss). To enumerate cell sub-populations in the jejunum, the number of lamina propria lymphocytes (LPL) and IEL were counted per 300 μm length of villi and per 150 x 150 μm^2^ area of crypts. Cells were counted from 3 different areas per section and 3 villi or crypt regions were selected per area.

### Statistical analysis

All statistical analysis was conducted with Minitab 17 software (Minitab Inc., USA). Data were analysed for normality using the Anderson-Darling test and significance assessed by the Mann-Whitney U test. The Spearman’s rank correlation coefficient was calculated to evaluate relationships between parasitaemia and host immune responses, and each cytokine in the serum and jejunum.

## Acknowledgments

We are thankful to the staff at the Bumstead Unit at the National Avian Research Facility (NARF) for the supply of the chickens used in this study and to the staff at the Biological Services Unit at the RVC for their assistance in the care of the animals. We are also grateful to Edinburgh Genomics for their assistance with the transcriptome analysis. Also with thanks to Professor John Hopkins and Dr Mitch Abrahamson for their advice throughout this research.

## Funding

This research was supported by the Biotechnology and Biological Sciences Research Council (BBSRC) grants BB/L502455/1 and BB/L004046/1, in partnership with Cobb-Vantress Inc. This work was also supported by the Biotechnology and Biological Sciences Research Council Institute Strategic Program Grant BBS/E/D/20231760 and BBS/E/D/20231762 to The Roslin Institute.

This project has also received funding from the European Union’s Horizon 2020 Programme for research, technological development and demonstration under the Grant Agreement no. 633184. This publication reflects the views only of the author, and not the European Commission (EC). The EC is not liable for any use that may be made of the information contained herein.

The funders had no role in study design, data collection and interpretation, or the decision to submit the work for publication.

## References

1. Williams RB, Carlyle WW, Bond DR, Brown IA. 1999. The efficacy and economic benefits of Paracox, a live attenuated anticoccidial vaccine, in commercial trials with standard broiler chickens in the United Kingdom. Int J Parasitol 29:341–355.

2. Dalloul RA, Lillehoj HS. 2006. Poultry coccidiosis: recent advancements in control measures and vaccine development. Expert Rev Vaccines 5:143–163.

3. Long PL, Joyner LP. 1984. Problems in the identification of species of *Eimeria*. J Protozool 31:535–541.

4. Clark EL, Macdonald SE, Thenmozhi V, Kundu K, Garg R, Kumar S, Ayoade S, Fornace KM, Jatau ID, Moftah A, Nolan MJ, Sudhakar NR, Adebambo AO, Lawal IA, Alvarez Zapata R, Awuni JA, Chapman HD, Karimuribo E, Mugasa CM, Namangala B, Rushton J, Suo X, Thangaraj K, Srinivasa Rao AS, Tewari AK, Banerjee PS, Dhinakar Raj G, Raman M, Tomley FM, Blake DP. 2016. Cryptic Eimeria genotypes are common across the southern but not northern hemisphere. Int J Parasitol 46:537–544.

5. Long PL, Millard BJ. 1976. Studies on site finding and site specificity of *Eimeria praecox*, *Eimeria maxima* and *Eimeria acervulina* in chickens. Parasitology 73:327–336.

6. Yadav A, Gupta SK. 2001. Study of resistance against some ionophores in *Eimeria tenella* field isolates. Vet Parasitol 102:69–75.

7. Blake DP, Tomley FM. 2014. Securing poultry production from the ever-present *Eimeria* challenge. Trends Parasitol 30:12–19.

8. Bumstead JM, Bumstead N, Rothwell L, Tomley FM. 1995. Comparison of immune responses in inbred lines of chickens to *Eimeria maxima* and *Eimeria tenella*. Parasitology 111 (Pt 2):143–151.

9. Smith AL, Hesketh P, Archer A, Shirley MW. 2002. Antigenic diversity in *Eimeria maxima* and the influence of host genetics and immunization schedule on cross-protective immunity. Infect Immun 70:2472–2479.

10. Blake DP, Hesketh P, Archer A, Shirley MW, Smith AL. 2006. *Eimeria maxima*: the influence of host genotype on parasite reproduction as revealed by quantitative real-time PCR. Int J Parasitol 36:97–105.

11. Lillehoj HS, Ruff MD. 1987. Comparison of disease susceptibility and subclass-specific antibody response in SC and FP chickens experimentally inoculated with *Eimeria tenella*, E. acervulina, or E. maxima. Avian Dis 31:112–119.

12. Lillehoj HS. 1986. Immune response during coccidiosis in SC and FP chickens. I. In vitro assessment of T cell proliferation response to stage-specific parasite antigens. Vet Immunol Immunopathol 13:321–330.

13. Smith NC, Wallach M, Miller CM, Morgenstern R, Braun R, Eckert J. 1994. Maternal transmission of immunity to *Eimeria maxima*: enzyme-linked immunosorbent assay analysis of protective antibodies induced by infection. Infect Immun 62:1348–1357.

14. Davis PJ, Parry SH, Porter P. 1978. The role of secretory IgA in anti-coccidial immunity in the chicken. Immunology 34:879–888.

15. Yun CH, Lillehoj HS, Zhu J, Min W. 2000. Kinetic differences in intestinal and systemic interferon-gamma and antigen-specific antibodies in chickens experimentally infected with *Eimeria maxima*. Avian Dis 44:305–312.

16. Rose ME, Hesketh P. 1979. Immunity to coccidiosis: T-lymphocyte- or B-lymphocyte-deficient animals. Infect Immun 26:630–637.

17. Bumstead N, Millard B. 1987. Genetics of resistance to coccidiosis: response of inbred chicken lines to infection by *Eimeria tenella* and *Eimeria maxima*. Br Poult Sci 28:705–715.

18. Rothwell L, Gramzinski RA, Rose ME, Kaiser P. 1995. Avian coccidiosis: changes in intestinal lymphocyte populations associated with the development of immunity to *Eimeria maxima*. Parasite Immunol 17:525–533.

19. Hong YH, Lillehoj HS, Lillehoj EP, Lee SH. 2006. Changes in immune-related gene expression and intestinal lymphocyte subpopulations following *Eimeria maxima* infection of chickens. Vet Immunol Immunopathol 114:259–272.

20. Djeraba A, Bernardet N, Dambrine G, Quere P. 2000. Nitric oxide inhibits Marek’s disease virus replication but is not the single decisive factor in interferon-gamma-mediated viral inhibition. Virology 277:58–65.

21. Mallick AI, Haq K, Brisbin JT, Mian MF, Kulkarni RR, Sharif S. 2011. Assessment of bioactivity of a recombinant chicken interferon-gamma expressed using a baculovirus expression system. J Interferon Cytokine Res 31:493–500.

22. Kogut MH, Rothwell L, Kaiser P. 2005. IFN-gamma priming of chicken heterophils upregulates the expression of proinflammatory and Th1 cytokine mRNA following receptor-mediated phagocytosis of *Salmonella* enterica serovar enteritidis. J Interferon Cytokine Res 25:73–81.

23. Heriveau C, Dimier-Poisson I, Lowenthal J, Naciri M, Quere P. 2000. Inhibition of *Eimeria tenella* replication after recombinant IFN-gamma activation in chicken macrophages, fibroblasts and epithelial cells. Vet Parasitol 92:37–49.

24. Lillehoj HS, Choi KD. 1998. Recombinant chicken interferon-gamma-mediated inhibition of *Eimeria tenella* development *in vitro* and reduction of oocyst production and body weight loss following *Eimeria acervulina* challenge infection. Avian Dis 42:307–314.

25. Lowenthal JW, York JJ, O’Neil TE, Rhodes S, Prowse SJ, Strom DG, Digby MR. 1997. *In vivo* effects of chicken interferon-gamma during infection with *Eimeria*. J Interferon Cytokine Res 17:551–558.

26. Yun CH, Lillehoj HS, Choi KD. 2000. *Eimeria tenella* infection induces local gamma interferon production and intestinal lymphocyte subpopulation changes. Infect Immun 68:1282–1288.

27. Rothwell L, Young JR, Zoorob R, Whittaker CA, Hesketh P, Archer A, Smith AL, Kaiser P. 2004. Cloning and characterization of chicken IL-10 and its role in the immune response to *Eimeria maxima*. J Immunol 173:2675–2682.

28. Haritova AM, Stanilova SA. 2012. Enhanced expression of IL-10 in contrast to IL-12B mRNA in poultry with experimental coccidiosis. Exp Parasitol 132:378–382.

29. Wu Z, Hu T, Rothwell L, Vervelde L, Kaiser P, Boulton K, Nolan MJ, Tomley FM, Blake DP, Hume DA. 2016. Analysis of the function of IL-10 in chickens using specific neutralising antibodies and a sensitive capture ELISA. Dev Comp Immunol 63:206–212.

30. Arendt MK, Sand JM, Marcone TM, Cook ME. 2016. Interleukin-10 neutralizing antibody for detection of intestinal luminal levels and as a dietary additive in *Eimeria* challenged broiler chicks. Poult Sci 95:430–438.

31. Psifidi A, Hesketh P, Stebbings A, Archer A, Salmon N, Matika O, Banos G, Bumstead N, Smith A, Kaiser P. 2013. Identification of SNP markers for resistance to coccidiosis in chickens, p 16. Abstract from Proceedings of the 8th European Poultry Genetics Symposium. Venice, ITALY.

32. Boulton K, Nolan MJ, Wu Z, Riggio V, Matika O, Harman K, Hocking PM, Bumstead N, Hesketh P, Archer A, Bishop SC, Kaiser P, Tomley FM, Hume DA, Smith AL, Blake DP, Psifidi A. 2018. Dissecting the Genomic Architecture of Resistance to Eimeria maxima Parasitism in the Chicken. Front Genet 9:528.

33. Guo P, Thomas JD, Bruce MP, Hinton TM, Bean AG, Lowenthal JW. 2013. The chicken TH1 response: potential therapeutic applications of ChIFN-gamma. Dev Comp Immunol 41:389–396.

34. Staines K, Hunt LG, Young JR, Butter C. 2014. Evolution of an expanded mannose receptor gene family. PLoS One 9:e110330.

35. Gobel TW, Chen CL, Shrimpf J, Grossi CE, Bernot A, Bucy RP, Auffray C, Cooper MD. 1994. Characterization of avian natural killer cells and their intracellular CD3 protein complex. Eur J Immunol 24:1685–1691.

36. Vervelde L, Jeurissen SH. 1993. Postnatal development of intra-epithelial leukocytes in the chicken digestive tract: phenotypical characterization in situ. Cell Tissue Res 274:295–301.

37. Jansen CA, van de Haar PM, van Haarlem D, van Kooten P, de Wit S, van Eden W, Viertlbock BC, Gobel TW, Vervelde L. 2010. Identification of new populations of chicken natural killer (NK) cells. Dev Comp Immunol 34:759–767.

38. Lillehoj HS. 1989. Intestinal intraepithelial and splenic natural killer cell responses to eimerian infections in inbred chickens. Infect Immun 57:1879–1884.

39. Wakelin D, Rose ME, Hesketh P, Else KJ, Grencis RK. 1993. Immunity to coccidiosis: genetic influences on lymphocyte and cytokine responses to infection with Eimeria vermiformis in inbred mice. Parasite Immunol 15:11–19.

40. Choi KD, Lillehoj HS, Zalenga DS. 1999. Changes in local IFN-gamma and TGF-beta4 mRNA expression and intraepithelial lymphocytes following Eimeria acervulina infection. Vet Immunol Immunopathol 71:263–275.

41. Vervelde L, Vermeulen AN, Jeurissen SH. 1996. In situ characterization of leucocyte subpopulations after infection with Eimeria tenella in chickens. Parasite Immunol 18:247–256.

42. Couper KN, Blount DG, Riley EM. 2008. IL-10: the master regulator of immunity to infection. J Immunol 180:5771–5777.

43. Chan IH, Wu V, Bilardello M, Mar E, Oft M, Van Vlasselaer P, Mumm JB. 2015. The Potentiation of IFN-gamma and Induction of Cytotoxic Proteins by Pegylated IL-10 in Human CD8 T Cells. J Interferon Cytokine Res 35:948–955.

44. Sand JM, Arendt MK, Repasy A, Deniz G, Cook ME. 2016. Oral antibody to interleukin-10 reduces growth rate depression due to Eimeria spp. infection in broiler chickens. Poult Sci 95:439–446.

45. Shanmugasundaram R, Sifri M, Selvaraj RK. 2013. Effect of yeast cell product (CitriStim) supplementation on broiler performance and intestinal immune cell parameters during an experimental coccidial infection. Poult Sci 92:358–363.

46. Cai CW, Blase JR, Zhang X, Eickhoff CS, Hoft DF. 2016. Th17 Cells Are More Protective Than Th1 Cells Against the Intracellular Parasite Trypanosoma cruzi. PLoS Pathog 12:e1005902.

47. Korn T, Bettelli E, Oukka M, Kuchroo VK. 2009. IL-17 and Th17 Cells. Annu Rev Immunol 27:485–517.

48. Noack M, Miossec P. 2014. Th17 and regulatory T cell balance in autoimmune and inflammatory diseases. Autoimmun Rev 13:668–677.

49. Kolls JK, Khader SA. 2010. The role of Th17 cytokines in primary mucosal immunity. Cytokine Growth Factor Rev 21:443–448.

50. Veldhoen M. 2017. Interleukin 17 is a chief orchestrator of immunity. Nat Immunol 18:612–621.

51. Kim WH, Jeong J, Park AR, Yim D, Kim YH, Kim KD, Chang HH, Lillehoj HS, Lee BH, Min W. 2012. Chicken IL-17F: identification and comparative expression analysis in Eimeria-infected chickens. Dev Comp Immunol 38:401–409.

52. Zhang L, Liu R, Song M, Hu Y, Pan B, Cai J, Wang M. 2013. Eimeria tenella: interleukin 17 contributes to host immunopathology in the gut during experimental infection. Exp Parasitol 133:121–130.

53. Del Cacho E, Gallego M, Lillehoj HS, Quilez J, Lillehoj EP, Ramo A, Sanchez-Acedo C. 2014. IL-17A regulates Eimeria tenella schizont maturation and migration in avian coccidiosis. Vet Res 45:25.

54. Skak K, Frederiksen KS, Lundsgaard D. 2008. Interleukin-21 activates human natural killer cells and modulates their surface receptor expression. Immunology 123:575–583.

55. Wendt K, Wilk E, Buyny S, Schmidt RE, Jacobs R. 2007. Interleukin-21 differentially affects human natural killer cell subsets. Immunology 122:486–495.

56. Perez-Mazliah D, Ng DH, Freitas do Rosario AP, McLaughlin S, Mastelic-Gavillet B, Sodenkamp J, Kushinga G, Langhorne J. 2015. Disruption of IL-21 signaling affects T cell-B cell interactions and abrogates protective humoral immunity to malaria. PLoS Pathog 11:e1004715.

57. Yi JS, Cox MA, Zajac AJ. 2010. Interleukin-21: a multifunctional regulator of immunity to infections. Microbes Infect 12:1111–1119.

58. Shi F, Erf GF. 2012. IFN-gamma, IL-21, and IL-10 co-expression in evolving autoimmune vitiligo lesions of Smyth line chickens. J Invest Dermatol 132:642–649.

59. Pot C, Jin H, Awasthi A, Liu SM, Lai CY, Madan R, Sharpe AH, Karp CL, Miaw SC, Ho IC, Kuchroo VK. 2009. Cutting edge: IL-27 induces the transcription factor c-Maf, cytokine IL-21, and the costimulatory receptor ICOS that coordinately act together to promote differentiation of IL-10-producing Tr1 cells. J Immunol 183:797–801.

60. Jeurissen SH, Janse EM, Vermeulen AN, Vervelde L. 1996. Eimeria tenella infections in chickens: aspects of host-parasite: interaction. Vet Immunol Immunopathol 54:231–238.

61. Wattrang E, Thebo P, Lunden A, Dalgaard TS. 2016. Monitoring of local CD8beta-expressing cell populations during Eimeria tenella infection of naive and immune chickens. Parasite Immunol 38:453–467.

62. Swinkels WJ, Post J, Cornelissen JB, Engel B, Boersma WJ, Rebel JM. 2007. Immune responses to an Eimeria acervulina infection in different broilers lines. Vet Immunol Immunopathol 117:26–34.

63. Jenkins MC, Augustine PC, Danforth HD, Barta JR. 1991. X-irradiation of Eimeria tenella oocysts provides direct evidence that sporozoite invasion and early schizont development induce a protective immune response(s). Infect Immun 59:4042–4048.

64. Vervelde L, Jeurissen SH. 1995. The role of intra-epithelial and lamina propria leucocytes during infection with Eimeria tenella. Adv Exp Med Biol 371B:953–958.

65. Norton CC, Joyner LP. 1975. The development of drug-resistant strains of Eimeria maxima in the laboratory. Parasitology 71:153–165.

66. Long PL, Millard BJ, Joyner LP, Norton CC. 1976. A guide to laboratory techniques used in the study and diagnosis of avian coccidiosis. Folia Vet Lat 6:201–217.

67. Johnson J, Reid WM. 1970. Anticoccidial drugs: lesion scoring techniques in battery and floor-pen experiments with chickens. Exp Parasitol 28:30–36.

68. Nolan MJ, Tomley FM, Kaiser P, Blake DP. 2015. Quantitative real-time PCR (qPCR) for Eimeria tenella replication--Implications for experimental refinement and animal welfare. Parasitol Int 64:464–470.

69. Li YP, Bang DD, Handberg KJ, Jorgensen PH, Zhang MF. 2005. Evaluation of the suitability of six host genes as internal control in real-time RT-PCR assays in chicken embryo cell cultures infected with infectious bursal disease virus. Vet Microbiol 110:155–165.

70. Sutton KM, Hu T, Wu Z, Siklodi B, Vervelde L, Kaiser P. 2015. The functions of the avian receptor activator of NF-kappaB ligand (RANKL) and its receptors, RANK and osteoprotegerin, are evolutionarily conserved. Dev Comp Immunol 51:170–184.

71. Kaiser P, Rothwell L, Galyov EE, Barrow PA, Burnside J, Wigley P. 2000. Differential cytokine expression in avian cells in response to invasion by Salmonella typhimurium, Salmonella enteritidis and Salmonella gallinarum. Microbiology 146 Pt 12:3217–3226.

72. Bolger AM, Lohse M, Usadel B. 2014. Trimmomatic: a flexible trimmer for Illumina sequence data. Bioinformatics 30:2114–2120.

73. Dobin A, Davis CA, Schlesinger F, Drenkow J, Zaleski C, Jha S, Batut P, Chaisson M, Gingeras TR. 2013. STAR: ultrafast universal RNA-seq aligner. Bioinformatics 29:15–21.

74. Liao Y, Smyth GK, Shi W. 2014. featureCounts: an efficient general purpose program for assigning sequence reads to genomic features. Bioinformatics 30:923–930.

75. Robinson MD, McCarthy DJ, Smyth GK. 2010. edgeR: a Bioconductor package for differential expression analysis of digital gene expression data. Bioinformatics 26:139–140.

76. Theocharidis A, van Dongen S, Enright AJ, Freeman TC. 2009. Network visualization and analysis of gene expression data using BioLayout Express(3D). Nat Protoc 4:1535–1550.

